# Guanosine-specific single-stranded ribonuclease effectors of a phytopathogenic fungus potentiate host immune responses

**DOI:** 10.1101/2021.10.13.464185

**Authors:** Naoyoshi Kumakura, Suthitar Singkaravanit-Ogawa, Pamela Gan, Ayako Tsushima, Nobuaki Ishihama, Shunsuke Watanabe, Mitsunori Seo, Shintaro Iwasaki, Mari Narusaka, Yoshihiro Narusaka, Yoshitaka Takano, Ken Shirasu

## Abstract

Plants activate immunity upon recognition of pathogen-associated molecular patterns. Although phytopathogens have evolved a set of effector proteins to counteract plant immunity, some effectors are perceived by hosts and induce immune responses. Here, we show that two secreted ribonuclease effectors, SRN1 and SRN2, encoded in a phytopathogenic fungus, *Colletotrichum orbiculare*, induce cell death in a signal peptide- and catalytic residue-dependent manner, when transiently expressed in *Nicotiana benthamiana*. The pervasive presence of *SRN* genes across *Colletotrichum* species suggested the conserved roles. Using a transient gene expression system in cucumber (*Cucumis sativus*), an original host of *C. orbiculare*, we show that SRN1 and SRN2 potentiate host pattern-triggered immunity. Consistent with this, *C. orbiculare* SRN1 and SRN2 deletion mutants exhibited increased virulence on the host. In vitro analysis revealed that SRN1 specifically cleaves single-stranded RNAs at guanosine, leaving a 3′-end phosphate. This activity has not been reported in plants. Importantly, the potentiation of *C. sativus* responses by SRN1 and SRN2 depends on the signal peptide and ribonuclease catalytic residues, suggesting that secreted SRNs cleave RNAs in apoplast and are detected by the host. We propose that the pathogen-derived apoplastic guanosine-specific single-stranded endoribonucleases lead to immunity potentiation in plants.

## Introduction

Plants and phytopathogens have developed mutual attack and defense systems over millions of years of coevolution. Plants are able to recognize pathogens through cell surface-localized pattern recognition receptors (PRRs). PRRs are able to perceive broadly conserved pathogen-associated molecular patterns (PAMPs), as well as damage-associated molecular patterns (DAMPs) that are host plant-derived molecules generated during pathogen invasion or cell damage. PAMPs and DAMPs include proteins, lipids, carbohydrates, and nucleic acids. Direct or indirect PAMPs/DAMPs perception by PRRs induce pattern triggered-immunity (PTI), which includes both local and systemic immune responses (Boutrot & Zipfel, 2017).

To counteract plant immune systems, pathogens have evolved secreted proteins, referred to as effectors, that inhibit host immune responses and allow pathogens to establish infection (Win *et al*., 2012). However, some effectors or their functions are recognized by host PRRs and induce immune responses. For example, the presence of Avr2, an effector protein of *Cladosporium fluvum*, is indirectly recognized by Cf-2, a PRR of tomato, and induces an immune response. Avr2 is secreted out from *C. fluvum* into the host apoplastic region, binds to Rcr3, a host plant-derived protease, and inhibits its enzymatic activity. Tomato indirectly senses Avr2 probably by detecting the modification of Rcr3 via Cf-2 and induces an immune response to inhibit *C. fluvum* infection (Dixon *et al*., 2000; Rooney *et al*., 2005; Tang *et al*., 2017). Thus, effectors can cause both positive and negative effects on the establishment of infection. However, the mechanistic diversity of effector recognition by host PRRs is largely unknown.

*Colletotrichum* species are fungal pathogens that cause anthracnose disease on a variety of plants including economically important crops, fruits, and vegetables (Dean *et al*., 2012; Cannon *et al*., 2012). Most *Colletotrichum* species adopt a hemibiotroph lifestyle, consisting of an early biotrophic phase with no visible symptoms and a later necrotrophic phase associated with host cell death. Due to the agricultural and scientific importance of *Colletotrichum* species, genome sequencing of these fungi has been performed (O’Connell *et al*., 2012; Gan *et al*., 2013, 2016, 2021; Baroncelli *et al*., 2014, 2016; Hacquard *et al*., 2016; Tsushima *et al*., 2019).

Several *Colletotrichum* effectors have been identified. For example, NIS1 from *Colletotrichum orbiculare*, a causal agent of Cucurbitaceae anthracnose disease, suppresses PTI by inhibiting plant immunity-related kinases (Yoshino *et al*., 2012; Irieda *et al*., 2019). For other instances, the homologous effectors CoDN3 from *C. orbiculare* and ChEC3 from *Colletotrichum higginsianum*, a pathogen that causes anthracnose disease on Brassicaceae plants, suppress plant cell death induced by NIS1 and NLP1, respectively, when they are expressed together in *N. benthamiana* (Yoshino *et al*., 2012; Kleemann *et al*., 2012). Recently, a highly conserved *Colletotrichum* effector candidate that induces host nuclear expansion and cell death was identified (Tsushima *et al*., 2021). However, *Colletotrichum* effectors such that induce an immune response in the host plant have not been reported.

Recently, there has been an increasing number of reports on RNAs in the apoplast, an interface of plant-fungal interactions. In *Arabidopsis thaliana*, apoplastic fluid contains diverse small and long-noncoding RNAs (Baldrich *et al*., 2019; Karimi *et al*., 2021). In addition, small RNAs are exchanged between host plants and colonizing organisms, such as parasitic plants or microbes (Weiberg *et al*., 2013; Wang *et al*., 2016; Zhang *et al*., 2016; Shahid *et al*., 2018; Cai *et al*., 2018). Thus, apoplast may serve a place of communications between the different organisms. However, the nature and roles of RNAs in the apoplast in plant-microbe interactions are still open questions.

Here, we show that *C. orbiculare* ribonuclease effectors potentiate host immune responses in their catalytic residue-dependent manner. By comparing the genomes of two different *Colletotrichum* species, we identified 21 conserved effector candidates that are expressed upon infection. Among these, secreted ribonuclease 1 (SRN1) and the close homolog SRN2 were found as cell death-inducing effectors when transiently expressed in *N. benthamiana*. Interestingly, however, neither SRN1 nor SRN2 induced cell death in *Cucumis sativus*, an original host of *C. orbiculare* from which the strain was isolated. Instead, SRN1 and SRN2 potentiated host immune responses; *C. orbiculare srn1 srn2* double deletion mutants showed increased virulence on *C. sativus*. Importantly, the potentiation of host immunity requires ribonuclease catalytic residues as well as the signal peptide, implying that SRNs cleave RNAs in the apoplast and are detected by the host. Consistent with this scenario, biochemical analysis revealed that SRN1 is a single-stranded RNA (ssRNA) specific ribonuclease, cleaving at guanosine and leaving a 3′-end phosphate. Collectively, our data suggest that the enzymatic nature of SRN1 secreted from a phytopathogenic fungus can be recognized by the host cell to drive plant immunity. Our study reveals a novel aspect of plant-microbe interaction mediated by specific single-stranded ribonuclease effectors.

## Materials and Methods

### Identification of conserved effector candidates

Conserved effector candidates among *C. orbiculare* and *C. higginsianum* were identified as described in Fig. 1a. Orthologs of *C. orbiculare* secreted proteins were identified in *C. higginsianum* (O’Connell *et al*., 2012) by performing a BLASTp search (E-value cut-off 1E-9). Hits were further filtered by searching for evidence of expression in *C. higginsianum* orthologs according to expression sequence tag (EST) data (Takahara *et al*., 2009), removing hits that were annotated with the keywords glycosylphosphatidylinositol (GPI), membrane, mitochondrial or cytochrome, and retaining sequences that encoded proteins of less than 350 amino acids. To identify genes with potential roles in infection, only the highest scoring BLASTp hits of *C. orbiculare* genes that had previously been shown to be up-regulated *in planta* (Gan *et al*., 2013) were selected (Supporting Information Table S1).

**Figure 1.**
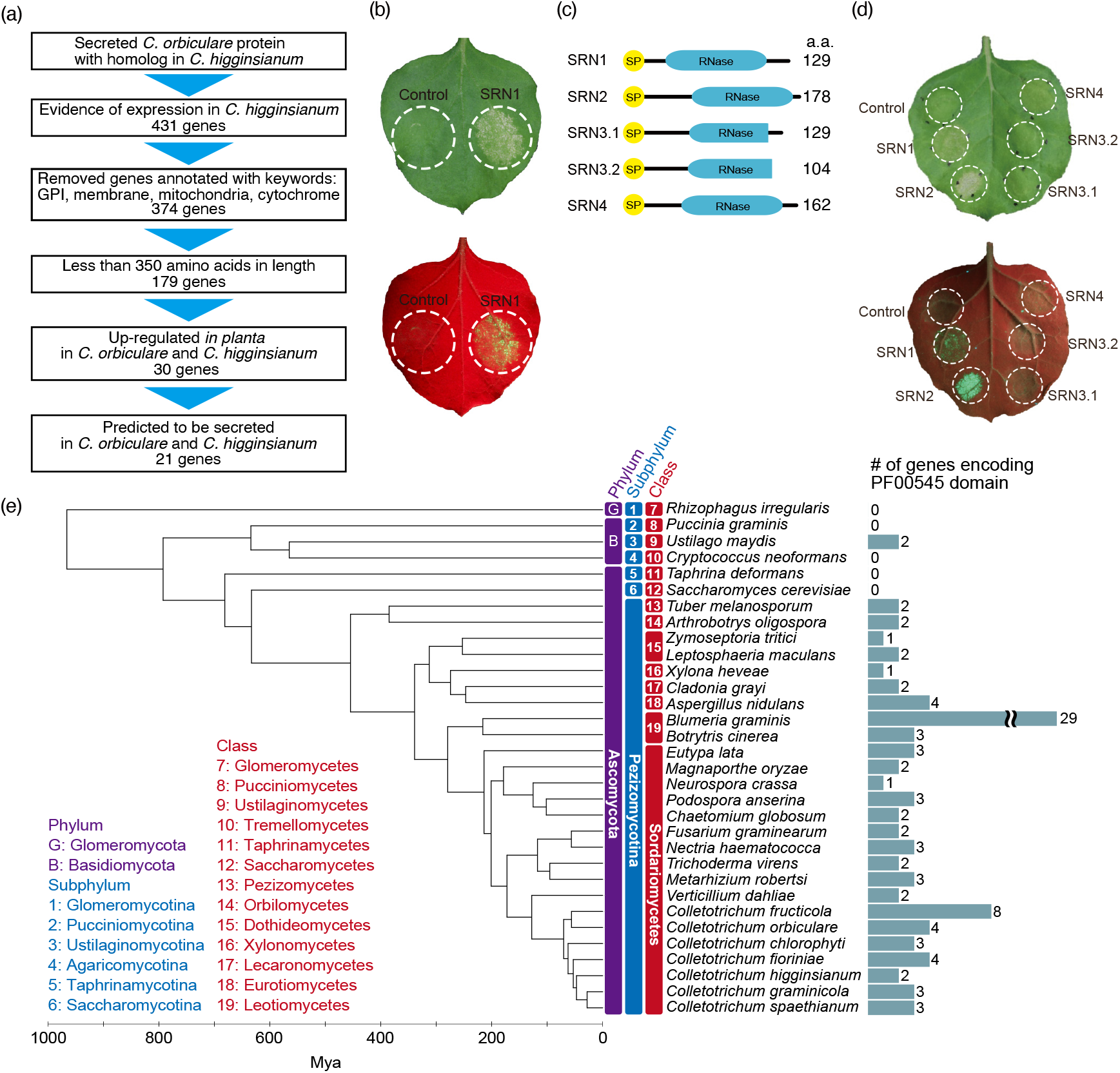
Identification of conserved *Colletotrichum* effectors that induce cell death in *N. benthamiana*. (a) Effector prediction pipeline in *C. orbiculare* and *C. higginsianum*. (b) *N. benthamiana* leaf expressing *C. orbiculare SRN1* using the *Agrobacterium*-mediated transient gene expression system. The pGWB2 binary vector was used. An *Agrobacterium* strain transformed with an empty vector was used for control. Photographs were taken at 6 days post-inoculation (dpi). The bottom image was taken under ultraviolet illumination to visualize plant cell death in green with autofluorescence. (c) Schematics of *C. orbiculare* SRN proteins. Yellow and blue boxes represent the signal peptides and RNase domains, respectively, that are conserved among SRN homologs. a. a. represents number of amino acids. (d) Cell death phenotypes of SRN1, SRN2, SRN3.1, SRN3.2, and SRN4 expressed as in (b). (e) PF00545 ribonuclease domain-containing proteins are highly conserved in fungi. Numbers of PF00545 ribonuclease domain-containing proteins in different fungi. Divergence dates were estimated based on a maximum likelihood tree constructed from 501 single copy genes in all the analyzed fungi using the program r8s.

### Identification of PF00545 ribonucleases from diverse fungi

Hmmscan was run against the Pfam 27.0 database using the default settings to identify proteins with the PF00545 ribonuclease domain (Finn *et al*., 2014; Eddy, 2011). Searches were run against 32 genomes from diverse fungi (Supporting Information Table S2) (Goffeau *et al*., 1996; Galagan *et al*., 2003; Loftus *et al*., 2005; Dean *et al*., 2005; Kämper *et al*., 2006; Cuomo *et al*., 2007; Espagne *et al*., 2008; Coleman *et al*., 2009; Martin *et al*., 2010; Spanu *et al*., 2010; Rouxel *et al*., 2011; Kubicek *et al*., 2011; Duplessis *et al*., 2011; Goodwin *et al*., 2011; Klosterman *et al*., 2011; Amselem *et al*., 2011; Yang *et al*., 2011; Berka *et al*., 2011; Arnaud *et al*., 2012; O’Connell *et al*., 2012; Gan *et al*., 2013, 2016, 2017; Blanco-Ulate *et al*., 2013; Cissé *et al*., 2013; Tisserant *et al*., 2013; Baroncelli *et al*., 2014; Hu *et al*., 2014; Gazis *et al*., 2016; Zampounis *et al*., 2016).

Proteins identified with the PF00545 domain were aligned by MAFFT and trimmed using trimAl (Katoh *et al*., 2002; Capella-Gutiérrez *et al*., 2009) using the default automated settings in both programs. The trimmed alignment was then used to construct a maximum likelihood tree with RAxML using the PROTAUTOGAMMA setting and 1,000 bootstrap replicates (Stamatakis, 2006). The conservation of active sites was assessed by checking for residues corresponding to *Aspergillus oryzae* ribonuclease T1 (RNase T1) Y64, H66, E84, R103, and H118 in the conserved domain cd00606 (NCBI’s conserved domain database) (Marchler-Bauer *et al*., 2017).

To generate the fungal species phylogenetic tree, single copy gene families were identified by orthoMCL (Li *et al*., 2003) from the 32 fungi analyzed using all-vs-all BLASTp with a cut-off E-value of 1E-5 and an inflation value of 1.5. Sequences from individual gene families were aligned using MAFFT and trimmed using trimAl (Katoh *et al*., 2002; Capella-Gutiérrez *et al*., 2009) as described above. Then, trimmed alignments from the 501 single copy gene families identified were concatenated resulting in a dataset of 227,412 sites. The concatenated alignment was used for RAxML analysis which was carried out as described above. *Rhizophagus irregularis* was set as the root using FigTree v1.4.2 (Rambaut, Andrew) in the best estimated tree, which was then converted to an ultrametric chronogram using r8s version 1.8 (Sanderson, 2003) using the Langley-Fitch molecular clock model. Previously estimated divergence times of 443–695 million years ago (mya) for Pezizomycotina-Saccharomycotina, 400-583 mya for the Pezizomycotina crown group, 267-430 mya for the Leotiomycetes-Sordariomycetes, 207-339 mya for Sordariomycetes, 487-773 mya for the Ascomycete crown (Beimforde *et al*., 2014) and 47 mya for the divergence between *C. graminicola* and *C. higginsianum* (O’Connell *et al*., 2012) were used to calibrate the tree. The phylogenetic trees generated were visualized in the interactive Tree of Life (Letunic & Bork, 2016).

### Prediction of SRN homologs in 22 *Colletotrichum* species

Using amino acid sequences of *C. orbiculare* SRN1, SRN2, SRN3.1, SRN3.2, and SRN4 as queries, genomes of 22 *Colletotrichum* species were searched by Exonerate version 2.2 software. Proteins lacking signal peptides predicted by SignalP 4.1 software (Petersen *et al*., 2011) were removed. The sequences were aligned using Molecular Evolutionary Genetics Analysis (Mega) Version 7.0 (Kumar *et al*., 2016). The phylogenetic tree of the SRN homologs was then drawn using the same software.

### Plant growth conditions

*N. benthamiana* plants were grown in a mixture of equal amounts of Supermix A (Sakata Seed Corp.) and vermiculite in 8 cm TO poly-pots (Tokai Agri System) under 16 h light:8 h dark conditions at 25 °C. *C. sativus* strain Suyo (Sakata Seed Corp.) plants were grown in the same soil mix and incubated under 10 h light: 14 h dark conditions at 24 °C.

### Plasmids

Plasmids used in this study are listed in Supporting Information Table S3. The method for plasmid construction is described in Supporting Information Method S1. Primers used in this study are listed in Supporting Information Table S4.

### Transient gene expression in *N. benthamiana* and *C. sativus*

*Agrobacterium*-mediated transient gene expression was performed following the previously described method with modifications (Chen *et al*., 2021). The *Agrobacterium tumefaciens* GV3101 and GV2260 strains were used. GV2260 strains were transformed with both pBBR*gabT* (Nonaka *et al*., 2017) and pEAQ-based plasmids (Sainsbury *et al*., 2009). The *Agrobacterium* culture was washed and resuspended in infiltration solution (10 mM MES pH 5.6, 10 mM MgCl_2_, and 150 µM acetosyringone). Infiltration solutions were at a density of O.D. 600 = 0.3. Each infiltration solution was infiltrated into 4-5 week-old *N. benthamiana* leaves or 6-9 day-old cotyledons of *C. sativus* using 1 ml syringes (TERUMO). An ultraviolet lamp MODEL B-100AP (UVP) was used for UV illumination. Photographs were taken using an EOS Kiss X6i (Canon). For UV illuminated leaves, a Y2 Professional Multi Coated Camera Lens Filter (Kenko) was used.

### RNA isolation, cDNA synthesis, and RT-qPCR

For obtaining vegetative hyphae (VH), *C. orbiculare* wild-type strain 104-T (MAFF 240422) was cultured on potato dextrose agar (PDA) medium (Nissui) then transferred onto potato dextrose (BD) liquid media and incubated for 3 days at 25 °C in the dark. For obtaining conidia, *C. orbiculare* hyphae were inoculated onto PDA media and incubated at 25 °C under black light blue light (10 h light: 14 h dark) for 6 days. Conidia were then suspended in water, filtered and collected by centrifugation. For one, three, and seven days post-inoculation (dpi), 1 × 10^6^ conidia ml^-1^ *C. orbiculare* conidia in 0.02% Silwet L-77 (Bio Medical Science) were inoculated onto the abaxial side of *C. sativus* cotyledons at 10 days post-germination (dpg) using a brush. Peeled epidermal cells were used for 1 and 3 dpi samples. Whole leaf tissues were used for 7 dpi samples. Three independent biological replicates were prepared for each sample.

For fungal biomass measurements, leaf disks were collected from *C. sativus* cotyledons infected with fungi at 88 hours post-inoculation (hpi) (as described in Fungal inoculation section) using a cork borer (4 mm diameter). Each sample consisted of at least six leaf disks from at least six leaves. Six replicates were prepared for each sample. All samples were transferred into 2-ml steel-top tubes, frozen using liquid nitrogen, and stored at -80 °C until RNA isolation. Total RNA isolation, DNA removal, cDNA synthesis and real-time quantitative PCR (RT-qPCR) reactions were performed as previously described (Kumakura *et al*., 2019) with slight modifications. First strand cDNA synthesis was performed with the ReverTra Ace qPCR RT Kit (TOYOBO) using the included primer mix, as well as gene-specific primers (listed in Supporting Information Table S4). Primer pairs used for RT-qPCR are also listed in Supporting Information Table S4.

### Fungal transformation and inoculation

The methods for fungal transformation and fungal inoculation are described in Supporting Information Method S2 and S3, respectively.

### Sequence alignment of SRNs

Amino acid sequences were aligned using CLC Genomics Workbench8 (CLC Bio). All coding sequences of *C. orbiculare* SRNs used in this report were cloned from *C. orbiculare* cDNAs and sequenced for verification.

### Measurement of oxidative burst from leaf disks

To detect chitin-induced reactive oxygen species (ROS) bursts, eight leaf disks were collected from *C. sativus* leaves using a cork borer (4 mm diameter) (Kai industries Co., Ltd.). Leaf disks were floated for more than 10 h on sterile water in 96-well microplates (655075, Greiner Bio-One), then the water was substituted by a solution containing 10 mg ml^-1^ horseradish peroxidase (Sigma), 1 µM L-012 (Wako), and 10 µM chitin heptose (Oligo Tech). Luminescence was measured for 30 min using a TriStar2 LB942 multi-plate reader (Berthold) (Kadota *et al*., 2018).

### Immunoblotting

The method for immunoblotting is described in Supporting Information Method S4.

### Protein deglycosylation enzyme treatment

Proteins isolated from *N. benthamiana* using the method described in the immunoblotting section were treated with Protein Deglycosylation Mix II (New England BioLabs) following the manufacturer’s protocol. Then, Samples were mixed with SDS-Laemmli buffer and analyzed by immunoblot.

### Recombinant protein expression and purification

The method for immunoblotting is described in Supporting Information Method S5.

### In vitro RNase assay

RNA substrates used in this study are shown in Fig. 6a, 6b, and Supplementary Information Fig. S7c. As ssRNA substrates, AG10 (5′-AGAGAGAGAGAGAGAGAGAG-3′), UC10 (5′-UCUCUCUCUCUCUCUCUCUC-3′), ssRNA1 (5 ’-AUCAUGCAUCAUCAUCAUCA-3 ’), ssRNA2 (5 ’-AUCAUCAUCAGUCAUCAUCA-3 ’), ssRNA3 (5 ’-AUCAUCAUCAUCAUCGAUCA-3 ’), ssRNA4 (5 ’-UCGCGUUGAUUACCCUGUUAUCCCUAGUGUACAU-3 ’) were chemically synthesized with fluorescein (FAM) addition at their 5′ end by Hokkaido System Science Co., Ltd. As a double-stranded RNA (dsRNA) substrate, dsRNA4 was prepared as following. ssRNA4 and chemically synthesized ssRNA (5 ’-AUGUACACUAGGGAUAACAGGGUAAUCAACGCGA-3 ’) which is complementary to ssRNA4 were mixed in buffer (20 mM Tris-HCl pH7.5, 150 mM NaCl, 1 mM DTT, and 2 mM MgCl_2_). The mixture was incubated at 95 °C for 5 min and then at room temperature for 30 min, resulting in dsRNA4.

For in vitro RNase assay, recombinant proteins or commercially available RNase T1 (Thermo Fisher Scientific) were mixed with 0.5 pmol substrate RNAs in 10 µl RNase reaction buffer with EDTA (20 mM Tris-HCl pH7.5, 150 mM NaCl, 1 mM DTT, and 5 mM EDTA) or 10 µl RNase reaction buffer with MgCl_2_ (20 mM Tris-HCl pH7.5, 150 mM NaCl, 1 mM DTT, and 5 mM MgCl_2_). The reaction was incubated for 30 min at 25 °C, then mixed with an equal volume of 2×RNA loading buffer [95% (v v^-1^) formamide, 0.025% (w v^-1^) SDS, and 0.5 mM EDTA], incubated 95 °C for 3 min, and cooled on ice for 1 min. The reaction was separated by 15% denaturing acrylamide urea gel electrophoresis. Signals of FAM labelled RNAs were detected using PharosFX (Bio-Rad) imaging systems.

### Linker ligation of RNAs

Linker ligation of RNAs was performed as previously described (Mito *et al*., 2020). Details are in Supporting Information Method S6.

## Results

### Prediction of *Colletotrichum* conserved effectors and identification of a cell death-inducing effector in *N. benthamiana*

To survey conserved effectors in the *Colletotrichum* genus, we reasoned that the candidates should be secreted to interact with host plant, conserved in the genus, and highly expressed during infection. Considering these criteria, we compared protein sequences from *C. orbiculare* and *C. higginsianum*, belonging to the orbiculare and destructivum species complexes, respectively. A total of 21 small secreted orthologous proteins, with evidence of up-regulation both in *C. higginsianum* and *C. orbiculare* during infection (Takahara *et al*., 2009; Gan *et al*., 2013), were selected as effector candidates conserved in the two pathogens (Fig. 1a, Supporting Information Table S1).

Among them, we found that one of the *C. orbiculare* effector candidates induced cell death (Fig. 1b) when expressed transiently in *N. benthamiana*. The gene that induced cell death (Locus tag: Cob_v010174) was named *<u>S</u>ECRETED <u>R</u>IBO<u>N</u>UCLEASE 1* (*SRN1*), due to the presence of the ribonuclease domain (Pfam database: PF00545, NCBI’s conserved domain database: cd00606) and the signal peptide sequence (Fig. 1c). The *C. orbiculare* genome is predicted to encode three other *SRN1-*like genes and we therefore termed them as *SRN2, SRN3* and *SRN4* (Supporting Information Fig. S1a, Table S6). By cloning the coding sequences from the cDNA of *C. orbiculare* we found that each gene transcript had a single isoform except for *SRN3*, which had two different isoforms (denoted as *SRN3*.*1* and *SRN3*.*2*) (Fig. 1c).

SRN1, SRN2, and SRN4 had five conserved ribonuclease catalytic residues inside their ribonuclease domains, while SRN3.1 and SRN3.2 contained only the first three (Supporting Information Fig. S1b) (Nishikawa *et al*., 1987; Noguchi *et al*., 1995; Marchler-Bauer *et al*., 2017).

To test the functional resemblance with *SRN1*, cell death induced by *SRN2, SRN3*.*1, SRN3*.*2*, and *SRN4* expressions was monitored in *N. benthamiana*. Like *SRN1, SRN2* induced cell death, while *SRN3*.*1, SRN3*.*2* and *SRN4* did not (Fig. 1d). Since the insufficient expression may hamper the conclusion, we harnessed the pEAQ-HT vector, which enables higher expression of proteins (Sainsbury *et al*., 2009). In this system, in addition to *SRN1* and *SRN2, SRN4* also induced cell death in *N. benthamiana* (Supporting Information Fig. S2a). The proteins with C-terminal HA tag did not impact on the cell death induced by *SRN1, SRN2*, and *SRN4* (Supporting Information Fig. S2b). We note that *SRN3*.*1-HA* weakly induced cell death in this high-expression system, while *SRN3*.*2-HA* did not, despite its expression being confirmed by immunoblot analysis (Supporting Information Fig. S2c).

### *SRN* homologs are conserved in all 22 *Colletotrichum* species tested

To analyze the conservation of *SRN1*, we surveyed the PF00545 ribonuclease domain in the Pfam database (Finn *et al*., 2014) because *SRN1* encodes the domain. Genes encoding PF00545 domains were conserved in bacteria and fungi, especially in Ascomycota, but not in plants and animals (Pfam 34.0) (Mistry *et al*., 2021) (Fig. 1e, 2), suggesting that PF00545 is the microorganisms specific domain. In fungi, all 26 Pezizomycotina species tested were predicted to encode proteins with the PF00545 domain. However, most species belonging to the Glomeromycota and Basidiomycota did not have the PF00545 domain, except for *Ustilago maydis*, a causal agent of corn smut.

**Figure 2.**
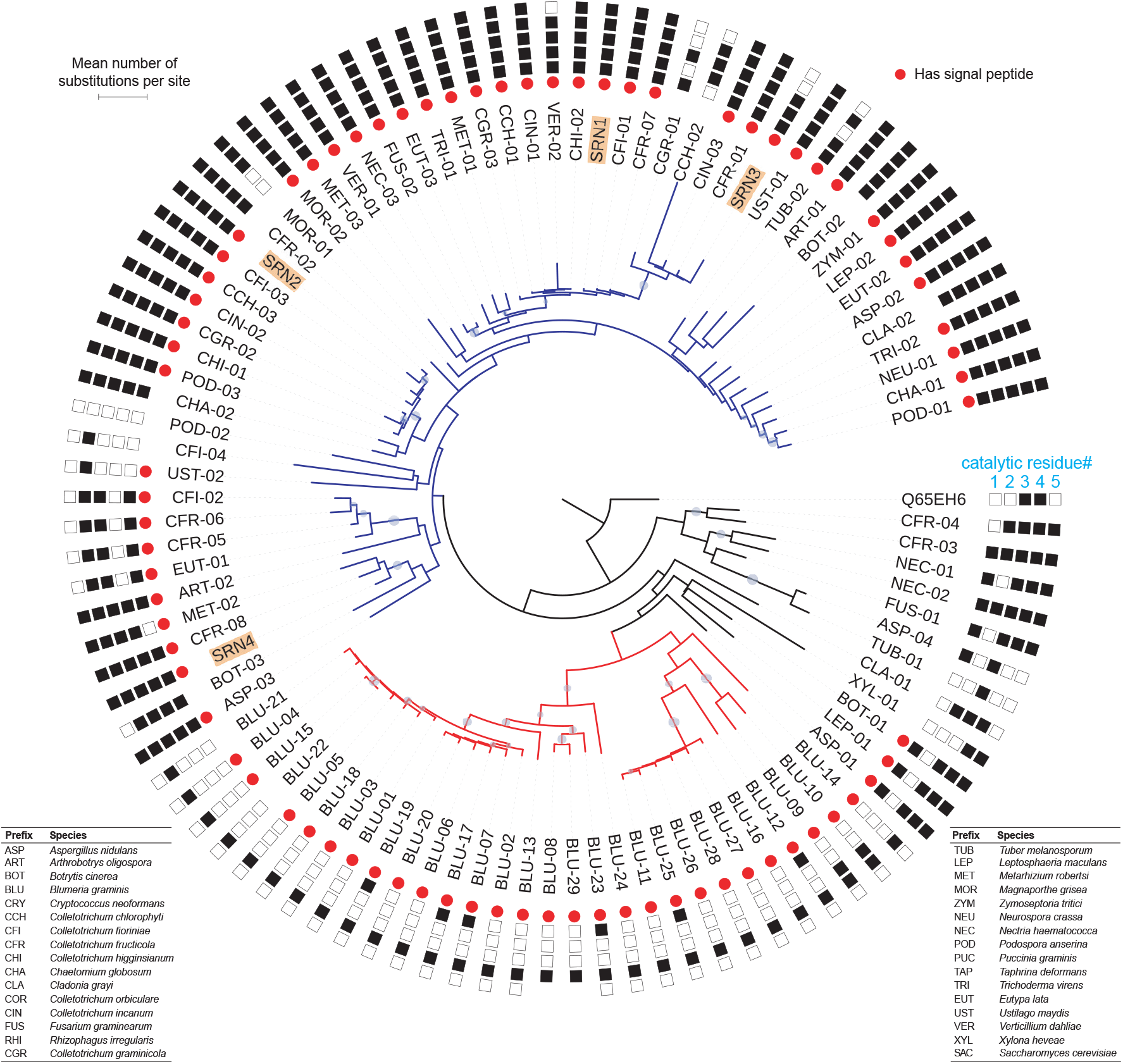
Phylogenetic relationship between fungal ribonuclease proteins. Maximum likelihood tree of sequences associated with the PF00545 ribonuclease domain in 32 different fungi drawn using the RAxML software. Grey circles on branches indicate branches with more than 50% bootstrap support values out of 1000 replicates. Black squares indicate conservation of five residues that are important for the ribonuclease catalytic activity. Red circles indicate the presence of a signal peptide according to the analysis by SignalP4.0.

To confirm if SRNs are conserved in the *Colletotrichum* genus, the genomes of 22 available *Colletotrichum* species (Supporting Information Table S7) were surveyed for the presence of SRN homolog-encoding sequences. Full-length amino acid sequences of *C. orbiculare* SRN1, SRN2, SRN3.1, SRN3.2, and SRN4 were used to query the whole genome sequences of the 22 species. All species tested had at least two SRN homologs (Supporting Information Table S7).

Based on the sequence similarity, the SRN homologs were classified into three groups named SRN1/3, SRN2, and SRN4. All 22 species had homologs belonging to the SRN1/3 and SRN2 groups, but only species from the gloeosporioides and orbiculare species complexes had the SRN4 group genes (Supporting Information Table S7). All species from the same species complex had the same composition of SRN homologs, except for the spaethianum species complex (Supporting Information Table S7).

### Cell death by *SRNs* is not observed in *C. sativus*, an original host of *C. orbiculare*

The SRN-mediated cell death observed in *N. benthamiana* led us to test the same phenotype in *C. sativus* (cucumber), an original host of *C. orbiculare*. To set out the protein expression in cucumber, we applied an *Agrobacterium*-mediated transient gene expression system established in melon (Chen *et al*., 2021) with several modifications. Here we used *A. tumefaciens* GV2260 strain which has the enhanced T-DNA translocation activity in certain plant species through expressing *gabT* gene (Nonaka *et al*., 2017). Indeed, this system allowed to accumulate GFP protein (as a marker protein) in fluorescence imaging (Supporting Information Fig. S3a) and in immunoblot (Supporting Information Fig. S3b). Harnessing this setup, we expressed *C. orbicualre* SRNs in *C. sativus*. Contrary to our expectation by *N. benthamiana* experiments, none of the *SRN* constructs induced detectable cell death on *C. sativus* cotyledons despite the detection of protein expression from these constructs (Fig. 3a, b).

**Figure 3.**
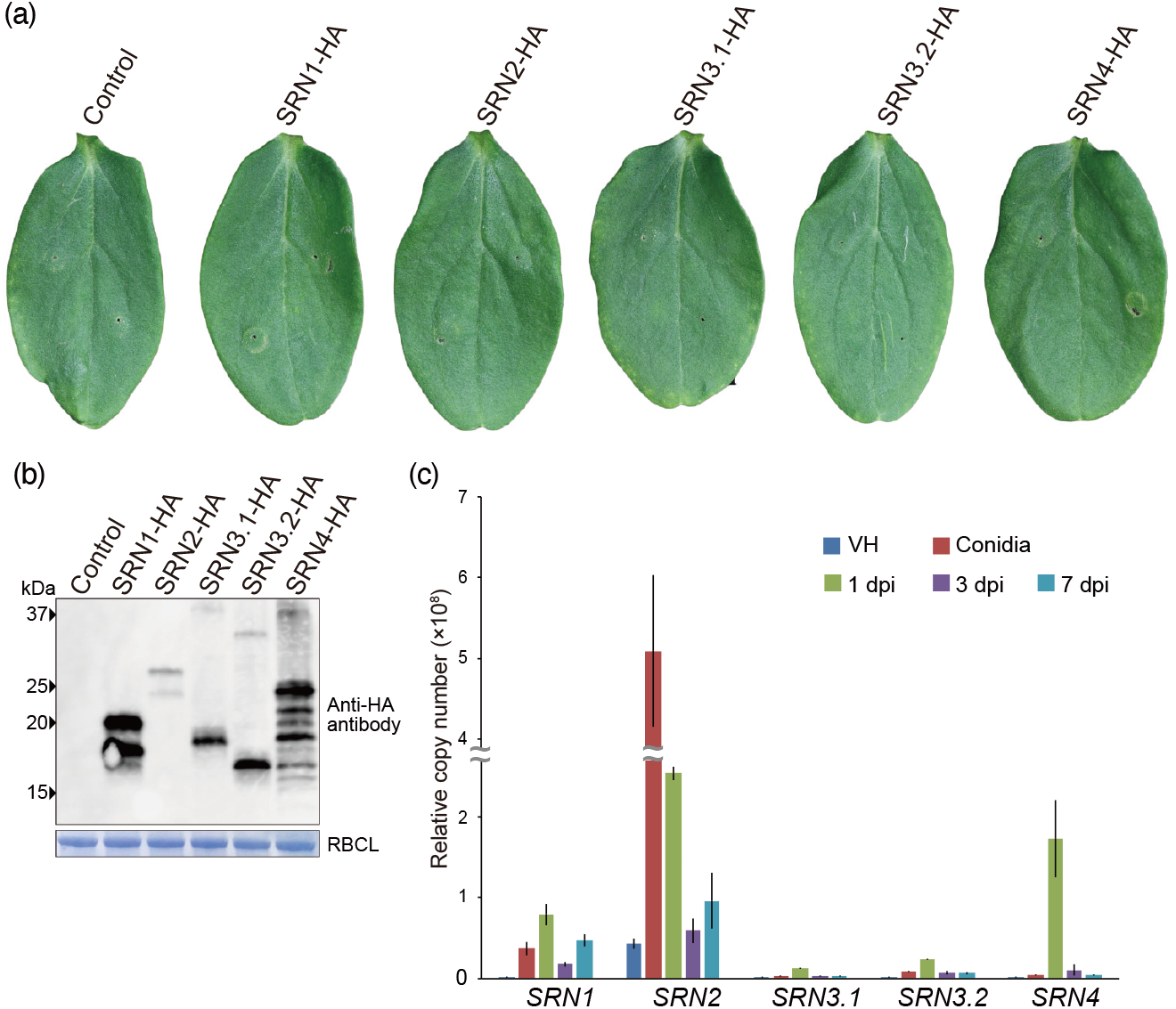
Expression of *C. orbiculare* SRNs on *C. sativus* leaves. (a) *C. sativus* cotyledons expressing HA-tagged SRNs using the *Agrobacterium*-mediated transient gene expression system. Photographs were taken at 5 dpi. Suspensions of *Agrobacteria* were infiltrated throughout the leaves. (b) Immunoblot analysis of proteins isolated from *C. sativus* cotyledons expressing SRNs. Total proteins were extracted at 5 dpi. The estimated molecular weight of each protein is as follows; SRN1-HA: 17.1 kDa, SRN2-HA: 22.9 kDa, SRN3.1-HA: 17.5 kDa, SRN3.2-HA: 14.8 kDa, SRN4-HA: 21.9 kDa. Anti-HA antibody (Roche) was used to detect tagged proteins. Coomassie-stained Rubisco large subunit (RBCL) proteins were used as loading controls. (c) Levels of *SRNs* transcripts during different infection stages of *C. orbiculare* were quantified using RT-qPCR. Total RNA was isolated from vegetative hyphae (VH) grown in vitro, conidia, and infected *C. sativus* leaves at 1, 3, and 7 dpi. To compare the number of transcripts from each gene, the copy number of each transcript was calculated using the standard curve drawn for the plasmid harboring the sequence of each transcript. Copy numbers were relative to the constitutively expressed *C. orbiculare* ribosomal protein L5 gene (Cob_v012718). Three biological replicates and two technical replicates were analyzed. Data represent mean ±SE.

### SRN1 and SRN2 enhance chitin-triggered ROS bursts in *C. sativus*

Given the tolerance to ectopic SRN expression in *C. sativus*, we were intrigued by the *SRN* genes expression profiles during *C. orbiculare* infection. RT-qPCR analysis revealed that all *SRNs*, except for SRN3.1 and SRN3.2, were strongly induced during infection compared to VH, non-inoculated fungal cells, especially at 1 dpi (Fig. 3c), implying that these effectors are likely to be involved in plant-fungi interaction at the early biotrophic phase. The upregulation of *SRNs* at 1 dpi prompted us to test if *SRNs* impact on plant immune responses in *C. sativus*. For this purpose, we monitored ROS bursts, a typical PTI response, triggered by chitin treatment (Torres *et al*., 2006). As shown in Supporting Information Fig. S4, chitin treatment strongly induced ROS bursts in *C. sativus* cotyledons.

Next, we examined the chitin-triggered ROS bursts in *C. sativus* cotyledons expressing either SRN1-HA, SRN2-HA, SRN3.1-HA, SRN3.2-HA, or SRN4-HA (Fig. 4a). Remarkably, *SRN1-HA* and *SRN2-HA* significantly enhanced chitin-triggered ROS bursts compared to GFP controls. On the other hand, SRN3.1-HA, SRN3.2-HA, and SRN4-HA did not, despite the detectable proteins accumulated (Fig. 4b). These data indicated that SRN1 and SRN2 leads to immune responses in host plants.

**Figure 4.**
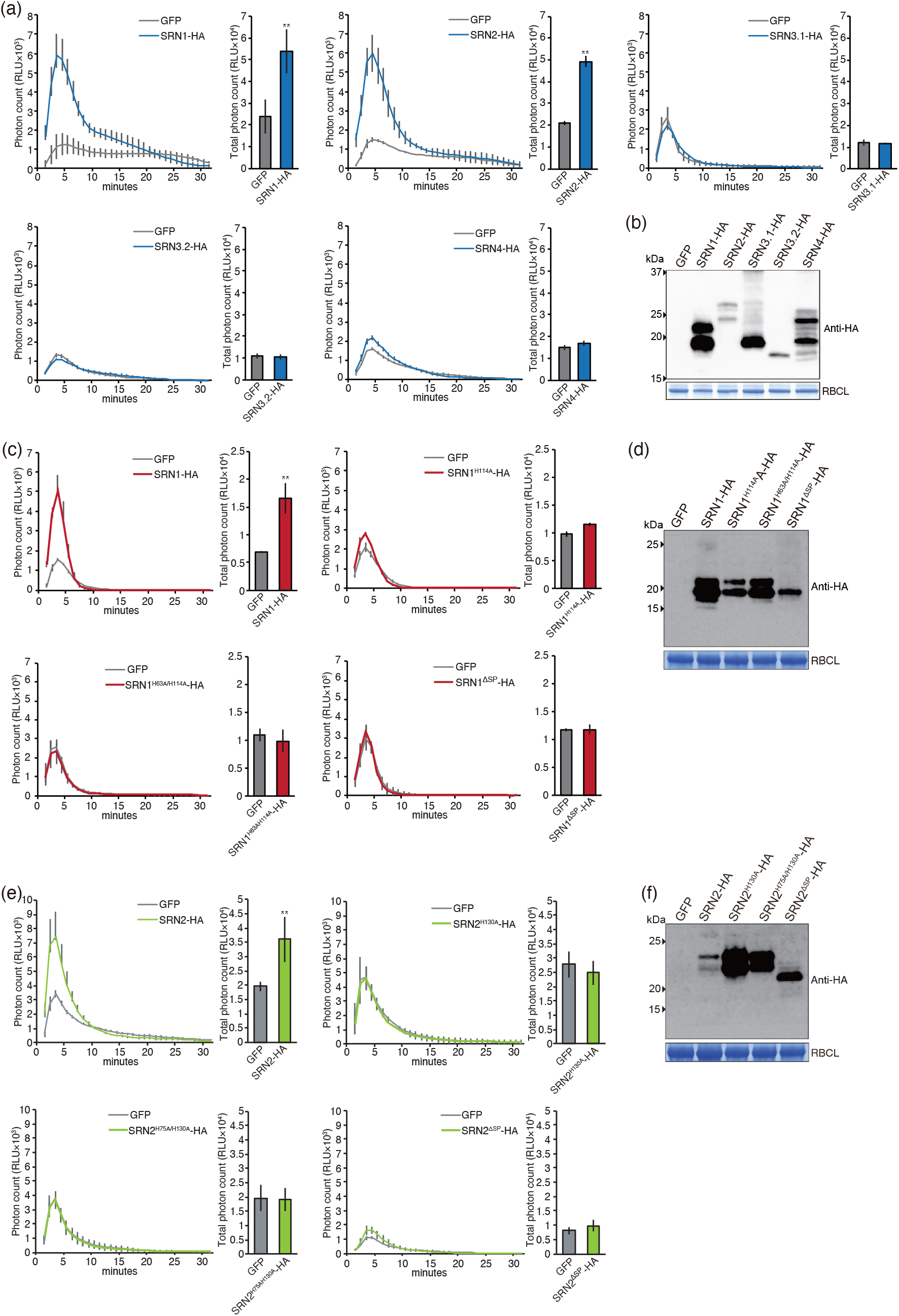
SRN1 and SRN2 expression potentiates chitin-triggered ROS bursts in *C. sativus* leaves. (a) Enhancement of chitin-triggered ROS bursts was observed in *C. sativus* cotyledons expressing SRN1-HA and SRN2-HA. Oxidative bursts were elicited by chitin (10 µM). Total photon counts were the sum of RLUs (relative light units) for a 30 min measurement. Three independent experiments showed similar results. Data represent mean ±SE (n=3). (b) Expression of SRN1-HA, SRN2-HA, SRN3.1-HA, SRN3.2-HA, and SRN4-HA proteins in *C. sativus* was confirmed by immunoblot analysis. Details of the analysis are the same as for the immunoblot in Fig. 3b. (c) Ribonuclease catalytic residues and the signal peptide of SRN1 are required for the chitin-triggered ROS burst enhancement. The H114 ribonuclease catalytic residue of SRN1 was mutated in SRN1^H114A^-HA. Both H63 and H114 ribonuclease catalytic residues of SRN1 were mutated in SRN1^H63A/H114A^-HA. The predicted signal peptide of SRN1 was deleted in SRN1^ΔSP^-HA. Experiments were performed as described in (a). Data represent mean ±SE (n=3). (d) Protein expression from the wild-type and mutated series of SRN1 (SRN1-HA, SRN1^H114A^-HA, SRN1^H63A/H114A^-HA, and SRN1^ΔSP^-HA) was confirmed by immunoblot analysis. The estimated molecular weight of each protein is as follows; SRN1-HA, SRN1^H114A^-HA and SRN1^H63A/H114A^-HA: 17.1 kDa, SRN1^ΔSP^-HA: 15.5 kDa. (e) Ribonuclease catalytic residues and the signal peptide of SRN2 were required for the chitin-triggered ROS burst enhancement. The H130 ribonuclease catalytic residue of SRN2 was mutated in SRN2^H130A^-HA. Both H75 and H130 ribonuclease catalytic residues of SRN2 were mutated in SRN2^H75A/H130A^-HA. The predicted signal peptide of SRN2 was deleted in SRN2^ΔSP^-HA. Experiments were performed as described in (a). Data represent mean ±SE (n=3). (f) Expression of wild-type and mutated series of SRN2 (SRN2-HA, SRN2^H130A^-HA, SRN2^H75A/H130A^-HA, and SRN2^ΔSP^-HA) was confirmed by immunoblot analysis. The estimated molecular weight of each protein is as follows; SRN2-HA, SRN2^H130A^-HA, and SRN2^H75A/H130A^-HA: 22.9 kDa, SRN2^ΔSP^-HA: 21.2 kDa. ^**^ indicates p < 0.01 (t-test) (a, c, e). Anti-HA antibody was used to detect HA-tagged proteins (b, d, f). Coomassie-stained RBCL proteins were used as loading controls (b, d, f).

### Enhancement of PTI by *SRN1*/*2* requires their catalytic residues and signal peptides

A previous report shows that two histidine residues (H40 and H92) are required for the ribonuclease catalytic activity of *A. oryzae* RNase T1, a homolog of SRNs (Nishikawa *et al*., 1987). To test whether histidine-dependent ribonuclease catalytic activity is involved in the ROS burst enhancement, we mutated the histidine residues of SRN1 corresponding to those of *A. oryzae*, H63 and H114, to alanine (SRN1^H63A/H114A^). Strikingly, the substitution abolished the SRN1-mediated ROS bursts (Fig. 4c). We also noticed that the single mutation of H114 alone reduced the enhancement of chitin-triggered ROS bursts (Fig. 4c). The similar double mutations (H92A and H130A) and single mutation (H130A) in SRN2 showed the same trends (Fig. 4e). Overall, we concluded that the ribonuclease catalytic residues of SRN1 and SRN2 are required for the enhancement of chitin-triggered ROS bursts.

Next, we assessed the effect of the signal peptide of SRN1 by deleting the sequence (SRN1^ΔSP^). The signal peptide-deleted SRN1 and SRN2 did not enhance chitin-triggered ROS bursts (Fig. 4c). These data suggest that the enhancement of the chitin-triggered response requires SRN1 and SRN2 to be external to the host cell, probably in the apoplastic region. We note that none of the loss of ROS burst enhancement by substitutions could be explained by the abrogation of protein expression (Fig. 4d and 4f).

The SRN proteins expressed in our setup were predicted to be modified post-translationally because all showed multiple bands larger than expected in immunoblots (Fig. 3b). Given that signal peptide-dependency for the mobility shift (Fig. 4d), one plausible post-translational modification is glycosylation.

Therefore, we predicted the glycosylation sites of SRNs using the NetNGlyc 1.0 server (Gupta & Brunak, 2002) and found that SRN1, SRN3.1, SRN3.2, and SRN4 have one potential glycosylated site each, while SRN2 has two. To assess whether glycosylation affects the function of SRNs, we mutated the predicted glycosylation sites (N101 and N143) of SRN2, as a representative of the SRNs, creating SRN2^N101Q/N143Q^ (Supporting Information Fig. S5a). As expected, this substitution constricted into a single protein band in immunoblot (Supporting Information Fig. S5b). Moreover, the treatment by deglycosylation enzyme reduced the intensity of the two bands in original SRN2 and generated a lower band, which was the same size as SRN2^N101Q/N143Q^ (Supporting Information Fig. S5c). However, irrespective of the glycosylation, we observed the immunity response by SRN2; SRN2^N101Q/N143Q^ induced chitin-triggered ROS bursts at the same level as the wild-type SRN2 in *C. sativus* (Supporting Information Fig. S5d-f) without any visible phenotype (Supporting Information Fig. S5g). In summary, these data suggest that the glycosylation of SRN2 does not affect its enhancement of chitin-triggered ROS bursts when expressed in *C. sativus*.

### Chitin-triggered MPK phosphorylation and PTI marker gene expression are enhanced by *SRN1* or *SRN2* in *C. sativus*

As the signaling pathways activated by PAMP perception include the activation of the mitogen-activated protein kinases, MPK3, MPK4, and MPK6 in *Arabidopsis* (Asai *et al*., 2002; Ichimura *et al*., 2006), we assessed the effect of SRN1 and SRN2 on chitin-triggered MPK phosphorylation in *C. sativus* (Fig. 5a, b). Indeed, SRN1 and SRN2 expression enhanced phosphorylation of MPKs (p44/42) upon chitin treatment, whereas the catalytically dead mutants (SRN1^H63A/H114A^ and SRN2^H75A/H130A^) did not.

**Figure 5.**
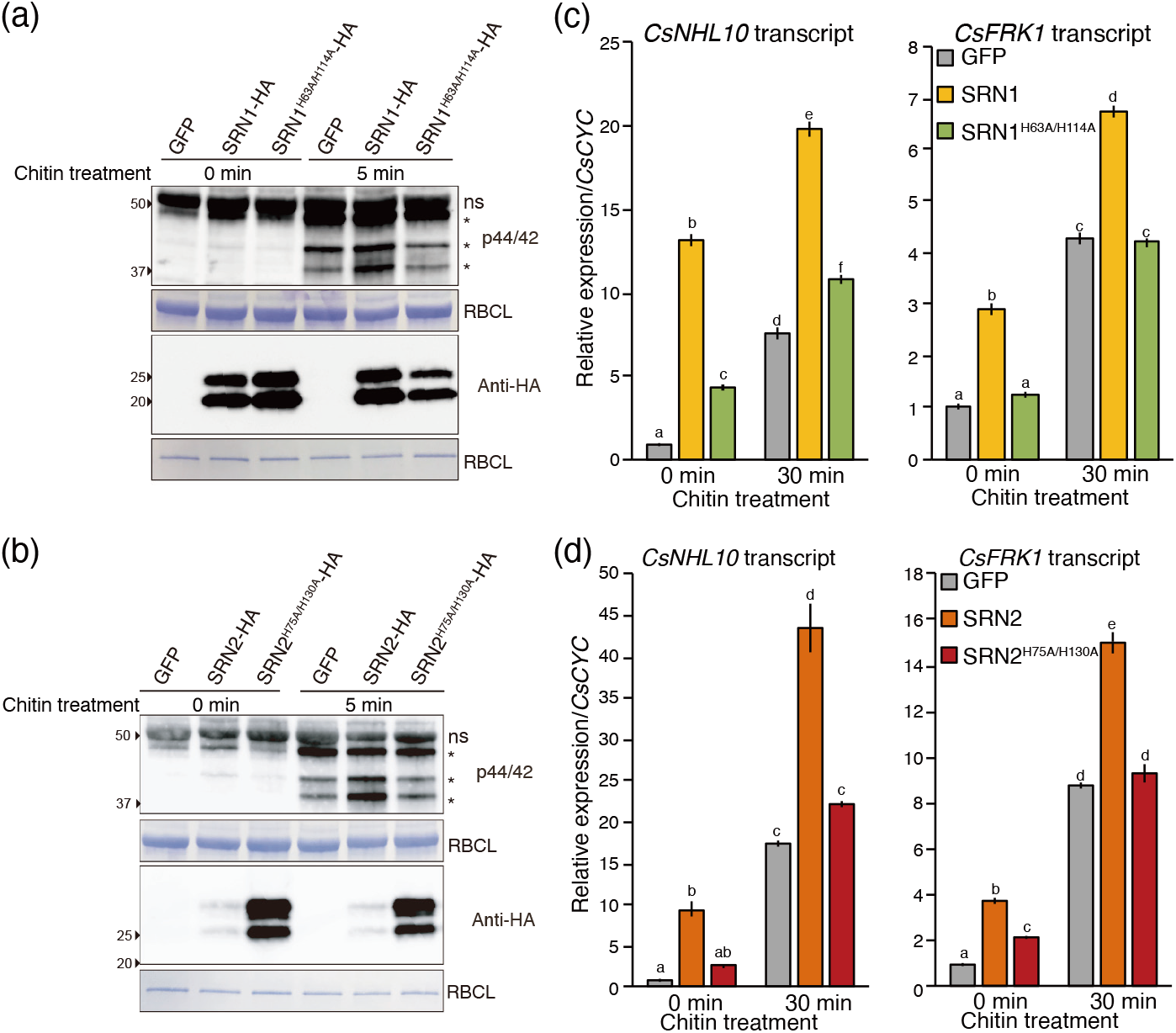
PTI potentiation by SRN1 and SRN2 expression in *C. sativus*. (a) Chitin-treated *C. sativus* cells expressing SRN1-HA showed enhanced MPKs phosphorylation. *C. sativus* cotyledons treated with chitin for 0 and 5 min were used. GFP, SRN1-HA, or SRN1^H63A/H114A^-HA were expressed in *C. sativus* cotyledons by an *Agrobacterium*-mediated transient gene expression system. Upper panel: phosphorylation of MPKs was detected using anti-phospho-p44/42 MAPK antibody. Lower panel: anti-HA antibody was used. Coomassie-stained RBCL proteins were used as loading controls. Similar results were obtained from independent three experiments. ns indicates a non-specific band. Asterisks indicate MPKs of *C. sativus*. (b) SRN1-HA expression induced PTI marker gene accumulation in *C. sativus* cells, as for SRN1-HA in panel (a). (b, d) Accumulation of *CsNHL10* and *CsFRK1* transcripts was quantified by RT-qPCR. *CsCYC* was used as endogenous control as established in a previous report (Liang *et al*., 2018). Primers used are listed in Supporting Information Table S3. Different lower-case letters indicate significant differences (p < 0.05, Tukey HSD). Data represent mean ±SE (n=3). Two independent experiments showed similar results. (c) Chitin-treated *C. sativus* cells expressing SRN2-HA showed enhanced MPKs phosphorylation. The experiment was performed as described in (a). (d) SRN2-HA expression induced PTI marker gene accumulation in *C. sativus* cells. The experiment was performed as described in (b).

It is well known that downstream events after activation of the MPK signaling cascade by PAMPs include transcriptional up-regulation of certain defense-related genes, such as *FRK1, NHL10, CYP82*, and *PHI1* in *Arabidopsis* (Wan *et al*., 2008). Therefore, we analyzed the expression of a set of *C. sativus* homologs of these PTI marker genes. We found that the expression of the *C. sativus FRK1* and *NHL10* homologs, *CsFRK1* and *CsNHL10*, was strongly induced 30 min after chitin treatment (Supporting Information Fig. S6). Importantly, these mRNAs were further induced by wild-type SRN1 and SRN2, but not by the catalytic mutants (Fig. 5c, d).

We note that even in the absence of chitin, SRN1 and SRN2 could lead *CsNHL10* and *CsFRK1* expression, suggesting the synergistic effects. Overall, these results indicates that SRN1 and SRN2 potentiate PTI responses in a catalytic residue-dependent manner.

### SRN1 cleaves ssRNAs at guanosine residues and leaves 3′-phosphates at 5′ fragments

To test an enzymatic activity of SRN1 in vitro, we utilized *Pichia pastoris*, which can secrete the recombinant protein outside the cell by secretion signal, α-factor (Brake *et al*., 1984). Here we substituted intrinsic signal peptide of SRN1 with the α-factor. Indeed, *P. pastoris* successfully expressed SRN1^ΔSP^ fused with α-factor at their N-terminus, however, the amount did not reach enough level for further biochemical use (data not shown), maybe due to cell toxicity. Therefore, a mutation, H63A, which could weaken the potential ribonuclease activity and cell toxicity was introduced into SRN1. H63 corresponds to H58 of *A. oryzae* RNase T1, and it has been reported that the ribonuclease activity is still detected and the substrate specificity is not affected when the mutation is introduced (Nishikawa *et al*., 1987). Indeed, sufficient amounts of SRN1^H63A/ΔSP^ protein were obtained for subsequent biochemical analyses. In addition to the single mutant, we isolated the double mutant (SRN1^H63A/H114A/ΔSP^) as a catalytically dead control.

Given that SRN1 homolog RNase T1 cleaves RNAs at guanosine residues specifically (Nishikawa *et al*., 1987), we reasoned that SRN1 may have the same nucleotide specificity. To test this possibility, we use two different ssRNAs as substrates: adenine/guanine-repeated polypurine RNA (AG10) and uracil/cytosine-repeated polypyrimidine RNA (UC10), conjugated with fluorescein (FAM) at their 5′ ends (Fig. 6a top). Indeed, SRN1^H63A/ΔSP^ digested AG10 but not UC10, indicating its nucleotide specificity toward. In contrast, the catalytically dead SRN1^H63A/H114A/ΔSP^ could not cleave the AG10 (Fig. 6a bottom). This reaction did not require Mg ions, which is a key cofactor for a subset of RNases (Fig. 6a, Supporting Information Fig. S7a). Titration of enzyme amount allowed us to track the reaction intermediates, which corresponds to nine fragments of AG10 (Supporting Information Fig. S7b), suggesting that SRN1 mediated endoribonucleolytic cleavage at adenosine or guanosine residues, but not both.

**Figure 6.**
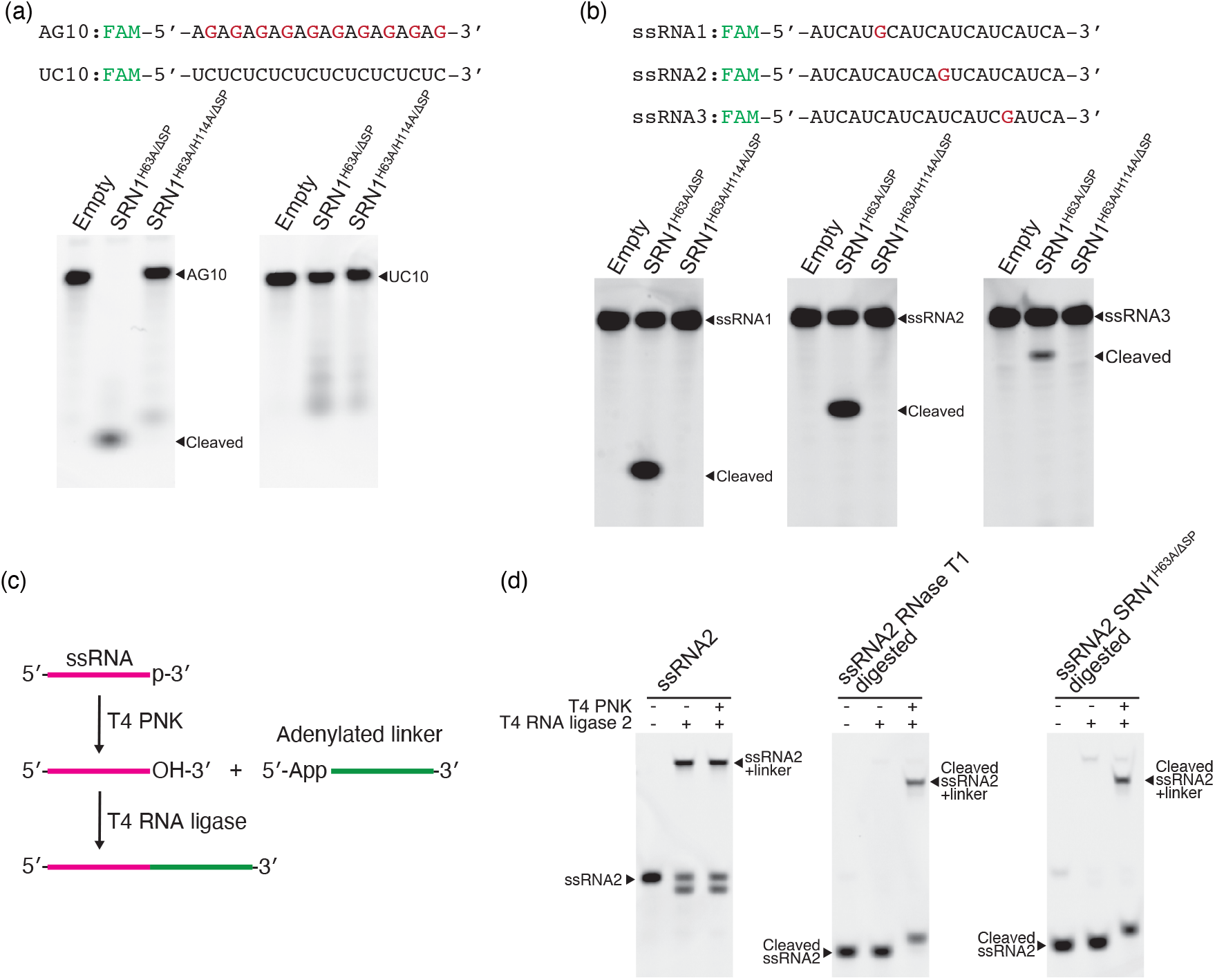
SRN1 cleaves ssRNAs at guanosine residues in vitro. (a) SRN1^H63A/ΔSP^ cleaved AG10, but did not UC10. In vitro RNase assays were performed with recombinant proteins produced by *P. pastoris* (SRN1^H63A/ΔSP^ and SRN1^H63A/H114A/ΔSP^), and chemically synthesized substrate ssRNAs (AG10 and UC10) labelled with fluorescein (FAM), at their 5′-termini. (b) SRN1^H63A/ΔSP^ cleaves ssRNAs at guanosine residues. ssRNA1, ssRNA2, and ssRNA3 have one guanosine residue, respectively, at different sites. ssRNA1, ssRNA2 and ssRNA3 are labelled with FAM at their 5′-termini. (c) Schematics of ssRNA dephosphorylation and linker ligation for (d). T4 PNK dephosphorylate 3′ end of ssRNA. T4 RNA ligase 2 conjugates the 3′-hydroxyl end of ssRNA with the pre-adenylated ssDNA linker. (d) 3′ end of cleaved ssRNA by SRN1^H63A/ΔSP^ possesses phosphate. ssRNA2 cleaved by SRN1^H63A/ΔSP^ was ligated with the pre-adenylated ssDNA only when the fragment was pre-treated with T4 PNK.

Thus, we tested whether SRN1 cleaves RNAs at guanosine residue using RNA substrates that only possess single guanosine but different positions (Fig. 6b, ssRNA1, ssRNA2, and ssRNA3). According to the position of the guanosine, SRN1^H63A/ΔSP^ generated short, middle, and long RNA fragments from ssRNA1, ssRNA2, and ssRNA3, respectively, showing that SRN1^H63A/ΔSP^ cleaves RNA at guanosine (Fig. 6b). We also tested whether SRN1 cleaves double stranded (ds) RNAs (Supplementary Information Fig. S7c, dsRNA4). Although un-annealed single strand RNA (ssRNA4) was cleaved by SRN1^H63A/ΔSP^ as expected, dsRNA4 was tolerate (Supplementary Information Fig. S7c). Thus, we concluded that SRN1 is a single-stranded RNA-specific endoribonuclease that cleaves at guanosine.

Considering that endonucleolytic cleavage by RNase T1 results in 3′-end phosphate at the 5′ RNA fragment (Nishikawa *et al*., 1987), we further investigated the molecular form of the cleaved end by SRN1. For this purpose, we harnessed the ligation-based assay; 3′ phosphate hampers the ligation to 3′ DNA fragment by T4 RNA ligase 2, whereas 3′ hydroxy group is susceptible to the reaction (Fig. 6c). As expected, the RNA (ssRNA2) that has a 3′-hydroxyl end was ligated (Fig. 6d, left panel). In contrast, the RNase T1-cleaved RNA could not engage in this ligation reaction unless the 3′ end is dephosphorylated by T4 polynucleotide kinase (PNK) (Fig. 6d, middle panel). Similarly, SRN1^H63A/ΔSP^ generated RNAs in the 3′-end form that could be ligated only after T4 PNK treatment (Fig. 6d, right panel). The striking correspondence of the substrate specificity (single stranded guanosine) (Fig. 6b), metal ion independency (Fig. 6a and Supporting Information Fig. S7a), and 3′-end phosphate in the cleaved product (Fig. 6d) showed that SRN1 functionally resembles RNase T1 (Nishikawa *et al*., 1987).

### The *srn1 srn2* double mutant strains show increased invasion and relative fungal biomass *in planta*

To assess the biological relevance of our findings, we established *srn1, srn2*, and *srn1 srn2* double knockout mutants and performed infection assays. We measured the invasion ratio, the percentage of successful infection hyphae per appressorium (Fig. 7a), at the early infection stage when the expression of *SRN1* and *SRN2* is induced (Fig. 3c). The invasion ratios were significantly increased in the two independent *srn1 srn2* double mutant strains (Fig. 7b).

**Figure 7.**
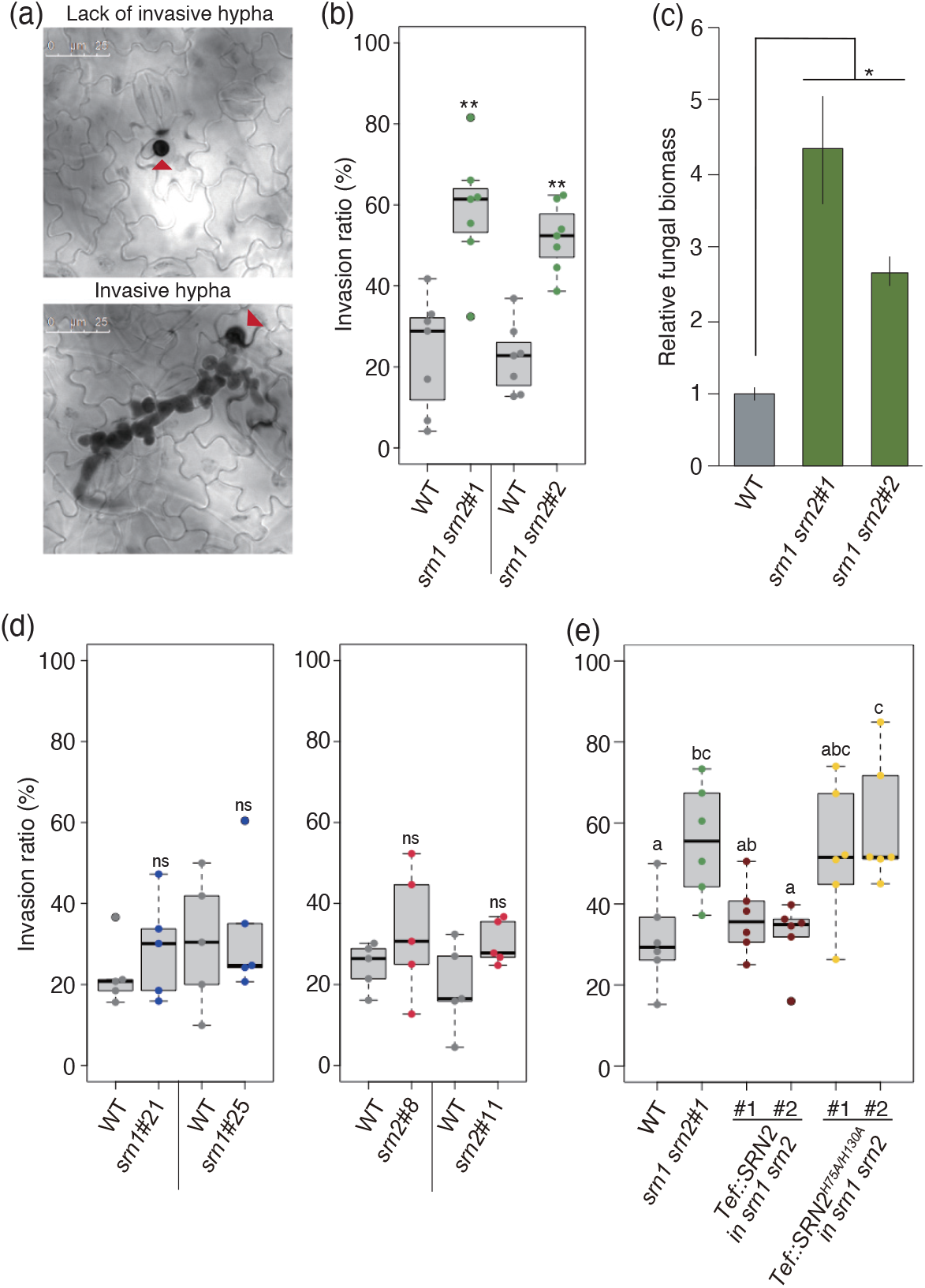
*C. orbiculare srn1 srn2* mutants showed increased virulence. (a) Example of lack of invasive hypha (top) and invasive hypha (bottom) formation from *C. orbiculare* appressoria. Red arrowheads indicate appressoria. Invasive hypha from successfully invaded appressoria were observed in the bottom panel. *C. sativus* cotyledons inoculated with *C. orbiculare* were trypan blue stained in both panels. Scale bars represent 25 µm. (b and d) Invasion ratio of a series of *srn* mutants on *C. sativus* cotyledons at 60 hpi. Leaves were inoculated with 5 µl of conidial suspensions at 1 × 10^5^ conidia ml^-1^. Each boxplot includes six to eight replicates. Each replicate was calculated using at least 50 appressoria. The box contains data within 1st and 3rd quartiles. ^*, **^, and ns indicate p < 0.05, p < 0.01, and not significant compared to *C. orbiculare* WT, respectively (t-test). (c) Fungal biomass during infection was quantified by RT-qPCR. A section of the ribosomal protein L5 transcript of *C. orbiculare* and a section of the *CsCYC* transcript of *C. sativus* were used for quantification. Primers used are listed in Supporting Information Table S4. Total RNAs were extracted from cotyledons inoculated either with *C. orbiculare* wild type or two *srn1 srn2* mutant strains at 88 hpi. ^*^ indicates p < 0.05 (t-test) compared to *C. orbiculare* WT. Data represent mean ±SE (n=6). (e) Overexpression of *SRN2* in *srn1 srn2* reduced the increased invasion ratio of the *srn1 srn2* mutant. *Tef* promoter-driven *SRN2* or *SRN2*^*H75A/H130A*^ were expressed in the *srn1 srn2*#1 mutant. The invasion ratio was measured using the same method as described in (b, d). Different lower-case letters indicate significant differences (p < 0.05, Tukey HSD).

In contrast, the *srn1* and *srn2* single mutants did not significantly alter the invasion ratios (Fig. 7d), suggesting the redundant functions of *SRN1* and *SRN2*.

To further ensure the role of *SRNs* in infection, we overexpressed *SRN2* in *srn1 srn2* double knockout strain. Here, ectopic *SRN2* was expressed by the promoter of *Aureobasidium pullulans TRANSLATION ELONGATION FACTOR* (*Tef*) (Wymelenberg *et al*., 1997). Indeed, the overexpression of SRN2 in the double knockout cells (denoted as *Tef::SRN2* in *srn1 srn2#1 and #2*) showed decreased invasion ratios compared to the parental double knockout mutant (*srn1 srn2*#1) (Fig. 7e). In contrast, catalytic inactive SRN2 (*Tef::SRN2*^*H75A/H130A*^ in *srn1 srn2*) could not complement the phenotype (Fig. 7e), suggesting the ribonuclease catalytic residues or activity are monitored by host to drive the immunity.

We also assessed the relative fungal biomass levels, which was probed by *C. orbiculare* transcripts (especially ribosome protein L5), during infection on *C. sativus* leaves. Consistent with the invasion ratios, *srn1 srn2* double mutants showed significantly increased fungal biomass (Fig. 7c). Collectively, our data indicates that *SRN1* and *SRN2* in *C. sativus* enhance defense responses of the host plant to this fungi.

## Discussion

Plants often perceive the presence of pathogens by recognizing molecules or the enzymatic activities of proteins originating from pathogens. Here, we report that *C. orbiculare* ribonuclease effectors, SRN1 and SRN2, potentiate typical PTI responses of *C. sativus* in a manner that is dependent on their catalytic residues and signal peptides. Our genetic analysis revealed that the *srn1 srn2* double mutants showed increased invasion ratios and relative fungal biomass (Fig. 7b, c), suggesting that SRN1 and SRN2 can be detrimental to the pathogen. Consistent with this notion, expression of SRN1 and SRN2 in *C. sativus* enhanced chitin-triggered ROS bursts (Fig. 4a) and MPK phosphorylation (Fig. 5a, b), as well as PTI marker gene expression (Fig. 5c, d). As these effects require catalytic residues and the signal peptides of SRN1 and SRN2, their enzymatic activity is likely to be recognized in the outside of the host cells (Fig. 4c, e), most probably in its apoplastic region. In line with this, in vitro analysis revealed that SRN1 recombinant proteins have an endoribonuclease activity that specifically cleaves ssRNAs at guanosine producing oligonucleotides with 3′ phosphate.

The action of SRN1 and SRN2 is apparently different from that of three ribonuclease-type effectors reported previously. Firstly, Zt6 from *Zymoseptoria tritici* is reported to be a host cell death-inducing effector by degrading rRNA in the host cells (Kettles *et al*., 2018). In contrast, SRN1 and SNR2 did not induce cell death in their host, *C. sativus*. In the *N. benthamiana* expression system, SRN2 induced cell death but their full cell death activity required signal peptide (Supporting Information Fig. S8), indicating that the effector targets are likely to be apoplastic RNAs, rather than cellular (r)RNA in the host. Secondly, CSEP0064/BEC1054, one of the 27 *Blumeria graminis* ribonuclease-like effectors that lack catalytic active residues, acts as a virulence factor inside the host cells (Pedersen *et al*., 2012; Pliego *et al*., 2013). More recently, Pennington *et al*. (2019) showed that CSEP0064/BEC1054 binds nucleic acids and inhibits the degradation of host rRNA induced by plant endogenous ribosome-inactivating proteins (RIPs) (Pennington *et al*., 2019). Based on these findings, Pennington *et al*. (2019) proposed that CSEP0064/BEC1054 is a pseudoenzyme that interacts with host ribosomes and inhibits the action of RIPs. Thirdly, AvrPm2, another ribonuclease-like protein of *B. graminis*, is recognized by the barley nucleotide-binding, leucine-rich repeat receptor (NLR) protein, Pm2 in the host cell (Praz *et al*., 2016), suggesting that AvrPm2 is a cytoplasmic effector that causes hypersensitive cell death triggered by an NLR. Thus, these three effectors are all predicted to be cytoplasmic effectors and are thus different from SRNs.

Host-specific cell death (only found in *N. benthamiana* but not in *C. sativus*) by SRNs expression remains an open question. One plausible explanation is that *N. benthamiana* encodes an as yet unidentified PRR that is able to trigger cell death upon direct or indirect detection of SRNs. Such cell death-inducing PRRs have been known in several species including potato and rice (Song *et al*., 1995; Du *et al*., 2015). In this scenario, it is also possible that *C. sativus* may encode a similar PRR that can detect the activity of *SRN1* and *SRN2* in the apoplast but potentiate PTI without causing cell death. As both signal peptides and catalytic residues of SRN1 and SRN2 are required for the full cell death activity in *N. benthamiana* and for PTI potentiation in *C. sativus*, it is possible that SRN1 and SRN2 cleaves RNAs in the apoplast and the resulting RNA molecules trigger immune responses via a PRR. In *Arabidopsis*, virus-derived dsRNAs induce PTI responses via SERK1, a receptor like kinase (Niehl *et al*., 2016). In addition, bacterial RNAs also induce immune responses in *Arabidopsis* when infiltrated into leaves (Lee *et al*., 2016). Thus, plants may be able to perceive certain RNA molecules in the apoplast. If RNAs are derived from host plants, these molecules may serve as DAMPs to indirectly detect invasive pathogens secreting specific RNases in the apoplast. This is a plausible case for SRNs, which are guanosine-specific single-strand endoribonucleases leaving 3′ phosphate, as plants normally do not encode RNases with this specificity. In mammals, PRRs such as Toll-like receptor (TLR) 3 and TLR7 can detect virus-derived dsRNA and ssRNAs, respectively (Takeuchi & Akira, 2010; Alexopoulou *et al*., 2001; Diebold & Brencicova, 2013). Isolation of such plant PRRs in the future will help to clarify the similarities and differences between the ways plants and animals recognize RNA molecules.

Why did all the *Colletotrichum* species we investigated encode SRN proteins? Although we did not detect a virulence function of SRN1 and SRN2 in our pathosystem, these proteins should provide biological advantage to the pathogen. For example, the function of these proteins is possibly manipulation of the local microbial community, as shown for Zt6 (Kettles *et al*., 2018; Snelders *et al*., 2018). Alternatively, SRNs target their own secreted RNAs. Fungal pathogens, such as *Botrytis cinerea*, can secrete small RNAs as effectors suppressing host immune responses (Weiberg *et al*., 2013). Thus, SRNs could be used to process such RNAs, which may serve as PAMPs when the host contains corresponding PRRs. However, if this is the case, it is difficult to explain why transient expression of SRNs in the host in the absence of a pathogen can induce immune responses. In addition, if SRNs are involved in the production of pathogen RNA effectors, the knockout phenotype is predicted to reduce virulence. However, the phenotype we observed was gain of virulence (Fig. 7). Another possibility is that SRNs target host apoplastic RNAs. *A. thaliana* apoplastic fluid contains both sRNAs and lncRNAs associated with proteins (Karimi *et al*., 2021). sRNAs of host plants were also detected in *Verticillium dahliae* and could down-regulate virulence-related genes (Zhang *et al*., 2016). Thus, such host-derived defensive apoplastic RNAs can be potential targets of SRNs. In this scenario, degrading host RNAs should increase pathogen virulence *per se*. Such virulence effects of SRNs may be observed in a host that is not able to detect the ribonuclease activity of SRNs. The identification of the target RNAs of SRNs will further clarify RNA-mediated plant-microbe interactions.

## Acknowledgements

We thank Drs. Satoko Nonaka and Hiroshi Ezura for providing the pBBR*gabT* plasmid, Drs. Mikiko Sodeoka and Kosuke Dodo for sharing equipment, and Dr. Yasuhiro Kadota for critical reading of the manuscript. This work was supported by Cooperative Research Grant of the Plant Transgenic Design Initiative (PTraD) by Gene Research Center, Tsukuba-Plant Innovation Research Center (University of Tsukuba), RIKEN Special Postdoctoral Researcher Program (NK), JSPS KAKENHI JP18K14440 and JP20K15500 (NK), JP19K15846 (PG), JP17J02983 (AT), JP21H05032 (YT), JP17H06172 (KS), JP19H02959 (SI), JST ACT-X Grant number JPMJAX20 (NK), The Program for Promotion of Basic and Applied Researches for Innovations in Bio-oriented Industry (BRAIN; 10103721) (KS), Council for Science, Technology and Innovation (CSTI) (KS), and Cross-ministerial Strategic Innovation Promotion Program (SIP) (YN, YT, and KS) “Technologies for creating next-generation agriculture, forestry and fisheries”. The authors declare no conflicts of interest associated with this manuscript.

## Author Contributions

NK, SO, YT, and KS designed the research. NK, SO, and MN performed experiments. PG and AT analyzed genome and transcriptome data. NK, NI, SW and MS prepared recombinant proteins. NK and SI performed in vitro analysis. YN, YT, and KS supervised the project. NK, PG, and KS wrote the manuscript with the edition from all the authors.

## Data Availability Statement

The data that supports the findings of this study are available in the supplementary material of this article or from the corresponding author upon reasonable request.

## Supporting Information

Figure S1. Alignment of *C. orbiculare* SRNs

Figure S2. SRNs expressed in *N. benthamiana* with the pEAQ binary vector

Figure S3. Establishment of the *Agrobacterium*-mediated transient gene expression system in *C. sativus*

Figure S4. Oxidative bursts are elicited by chitin on *C. sativus* cotyledons

Figure S5. The two predicted glycosylation sites in SRN2 are not involved in the potentiation of chitin-triggered ROS bursts

Figure S6. Assessment of PTI marker gene candidates in *C. sativus*

Figure S7. SRN1 does not cleave dsRNAs

Figure S8. Phenotype of *N. benthamiana* expressing SRN2 mutant proteins

Table S1. List of *Colletotorichum orbiculare* conserved effector candidates

Table S2. List of 32 fungal genomes

Table S3. Plasmid list

Table S4. DNA oligonucleotide list

Table S5. Fungal strain list

Table S6. Accession numbers of *SRN* genes

Table S7. Number of *SRN* genes in *Colletotrichum* species

Method S1. Plasmid construction

Method S2. Fungal transformation

Method S3. Fungal inoculation

Method S4. Immunoblotting

Method S5. Recombinant protein expression and purification

Method S6. Linker ligation of RNAs

## Notes

### Competing Interest Statement

The authors have declared no competing interest.

