## Supporting information for "Guanosine-specific single-stranded ribonuclease effectors of a phytopathogenic fungus potentiate host immune responses"

The following Supporting Information is available for this article:

**Fig. S1** Alignment of *C. orbiculare* SRNs
**Fig. S2** SRNs expressed in *N. benthamiana* with the pEAQ binary vector
**Fig. S3** Establishment of the *Agrobacterium*-mediated transient gene expression system in *C. sativus*
**Fig. S4** Oxidative bursts are elicited by chitin on *C. sativus* cotyledons
**Fig. S5** The two predicted glycosylation sites in SRN2 are not involved in the potentiation of chitin-triggered ROS bursts
**Fig. S6** Assessment of PTI marker gene candidates in *C. sativus*
**Fig. S7** SRN1 does not cleave dsRNAs
**Fig. S8** Phenotype of *N. benthamiana* expressing SRN2 mutant proteins

**Table S1** List of *Colletotorichum orbiculare* conserved effector candidates
**Table S2** List of 32 fungal genomes
**Table S3** Plasmid list
**Table S4** Oligonucleotide list
**Table S5** Fungal strain list
**Table S6** Accession numbers of *SRN* genes
**Table S7** Number of *SRN* genes in *Colletotrichum* species

**Method S1** Plasmid construction
**Method S2** Fungal transformation
**Method S3** Fungal inoculation
**Method S4** Immunoblotting
**Method S5** Recombinant protein expression and purification
**Method S6** Linker ligation of RNAs

**Fig. S1** Alignment of *C. orbiculare* SRNs

(a) Amino acid sequences of SRN1, SRN2, SRN3.1, SRN3.2, and SRN4. Predicted signal peptides and ribonuclease domains (cd00606) are highlighted in magenta and blue, respectively. Signal peptides were predicted by SignalP 4.1 server. (b) Amino acid sequences of SRN1, SRN2, SRN3.1, SRN3.2, and SRN4 are aligned and are shown using Mafft software and CLC Genomics Workbench software. Blue asterisks indicate putative ribonuclease catalytic residues corresponding to *Aspergillus oryzae* ribonuclease T1 Y64 (1st *), H66 (2nd *), E84 (3rd *), R103 (4th *), and H118 (5th *) in the conserved domain cd00606.

(a)

SRN1
MQFSVVLVSLFAIVASAVPTELQKRATTCGSTSYSTAQVNAASNAACQHVKAGTVAGTSTYPHRYNNYEGFDFDVTGPWYEFPIRTSGVYT
GGSPGADRVVINASCGQAGQITHTGASGNNFVGCSGTS

SRN2
MISLKSTLLLALCAFATASPLPEQTLQRRQGSSDNIQPVDCGGERYSRRQLEEATAEGCRLYANSLQVGTSQYPHRFNNREGLIFASTGPF
QEFPIIPSGNYSGRAPGPDRVVFDPDYRGRCSYVGAMTHTGADQRNGFVTCNETRRGGSGSSAAVSISPSGPLGVLLPLLSFMAFIA

SRN3.1
MFLGVAITAALSLEAHASPRHGYGRFTPRANDTEIVCGSTTYSLTEVRSAADAACQHVGAGTTAGSSTYPHPFRNSEGFEFHGVQEPFVEF
PMKTMGIYTGGKRLVRGCLASAFSEQIGDLASCADYHS

SRN3.2
MFLGVAITAALSLEAHASPRHGYGRFTPRANDTEIVCGSTTYSLTEVRSAADAACQHVGAGTTAGSSTYPHPFRNSEGFEFHGVQEPFVEF
PMKTMGIYTG

SRN4
MHSTKILVLLSASAASVLAVAVPPSDQNLQARAATVTCKPKTNLSTEKFVVNVDHAKAEAKKAGFTDGKSGYPHLFQNHDGIKWGVHNCDD
KKNPLQEYPVYWQGYKKQTVWNKEVKVKEQESTPLRVVYANKKGSIIYCGVMTHSKVEKNNRGLDYFQKCD

(b)

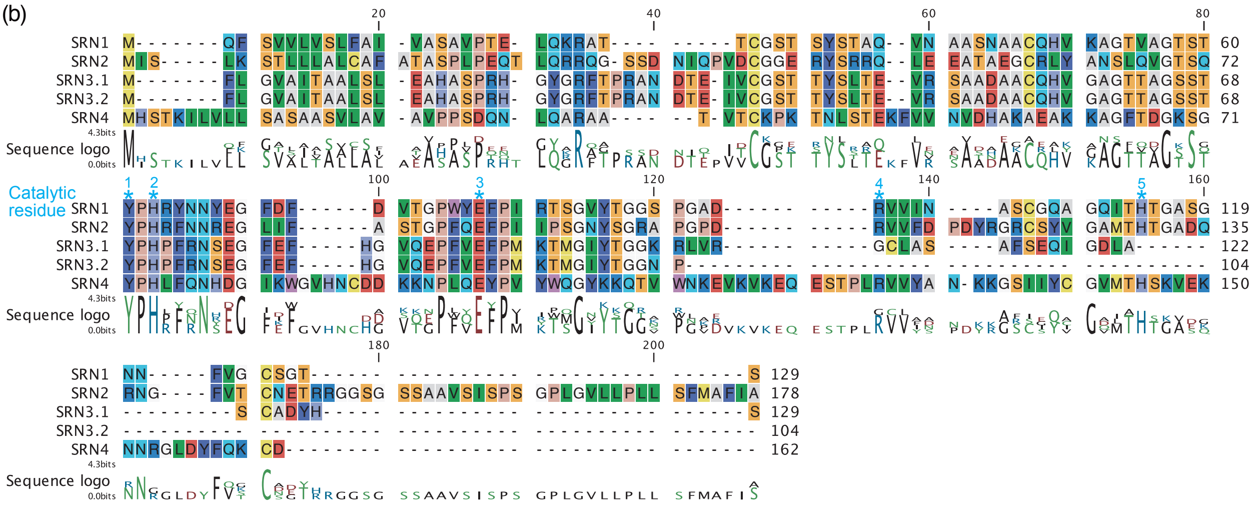

**Fig. S2** SRNs expressed in *N. benthamiana* with the pEAQ binary vector

(a) SRN1, SRN2, SRN3.1, SRN3.2 and SRN4 were expressed using an *Agrobacterium* mediated transient gene expression system. (b) SRN1-HA, SRN2-HA, SRN3.1-HA, SRN3.2-HA and SRN4-HA were expressed using the same method as (a). Photographs were taken at 5 dpi. The right hand photographs in (a, b) were taken under ultraviolet illumination. (c) Immunoblot analysis of proteins isolated from SRN-expressing *N. benthamiana*. Total proteins were extracted at 2 dpi. Estimated molecular weight of each protein is as follows; SRN1-HA: 17.1 kDa, SRN2-HA: 22.9 kDa, SRN3.1-HA: 17.5 kDa, SRN3.2-HA: 14.8 kDa, SRN4-HA: 21.9 kDa. Anti-HA antibody was used. Coomassie brilliant blue (CBB) stained Rubisco large subunit (RBCL) proteins were used as loading controls. pEAQ binary vector was used for expression. The backside of the same leaf as on the left was photographed under ultraviolet light, and the autofluorescence of cell death appears green in (a) and (b).

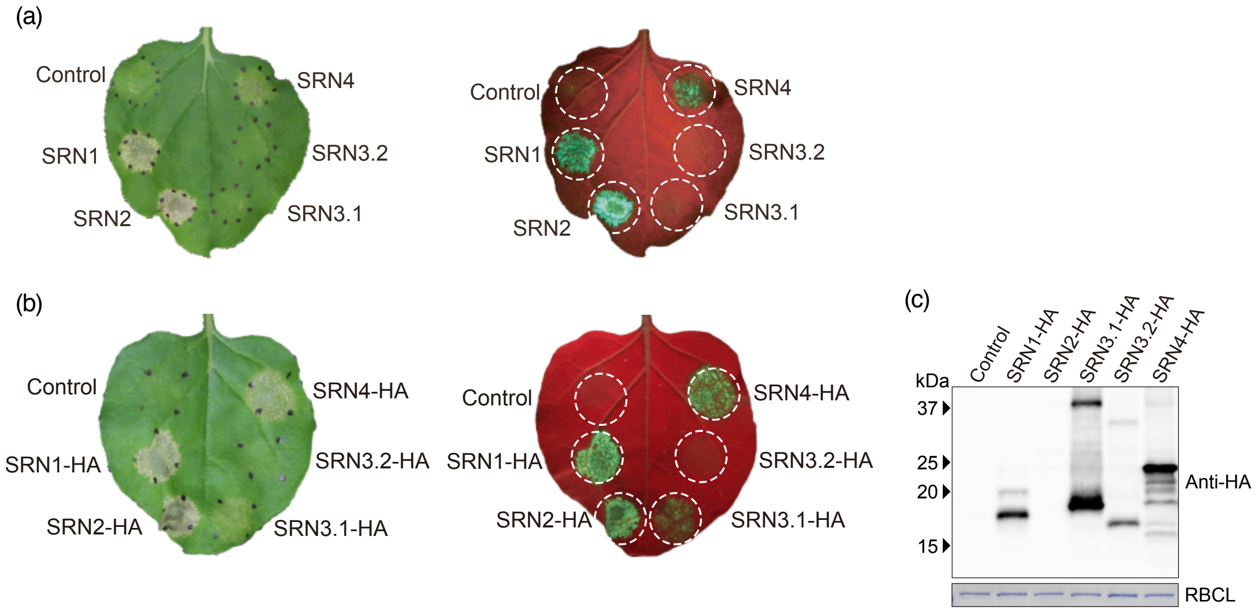

**Fig. S3** Establishment of the *Agrobacterium*-mediated transient gene expression system in *C. sativus*

(a) *C. sativus* cotyledon successfully expresses GFP using *A. tumefaciens* transformed with pEAQ-GFP and pBBR*gabT*. Photographs were taken at 6 dpi under ultraviolet illumination. (b) Immunoblot analysis of GFP protein. Total proteins were isolated at 6 dpi. Anti-GFP antibody was used for detection. CBB-stained RBCL proteins were used as loading controls.

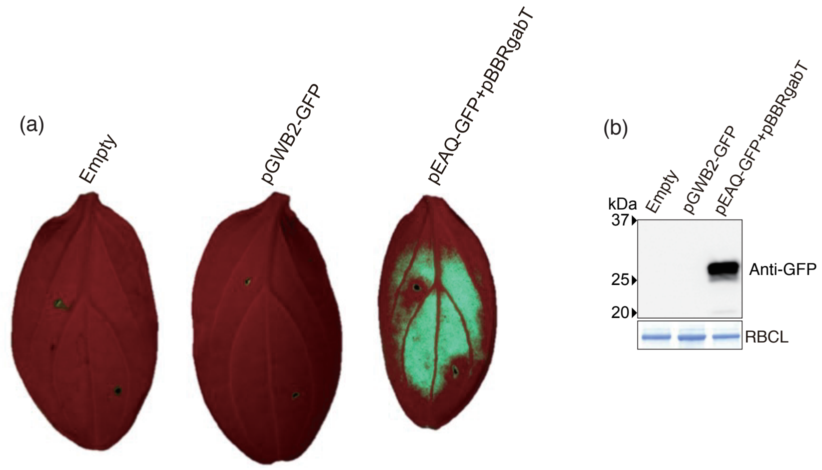


**Fig. S4** Oxidative bursts are elicited by chitin on *C. sativus* cotyledons

(a) Chitin treatment elicited oxidative burst on *C. sativus* cotyledons. Bars represent standard deviations. n=6. (b) Maximal and total ROS production after chitin treatment. n=6. Bars represent standard errors of six independent samples. * indicates $p < 0.05$ (t-test). Total photon counts were calculated using 20 times measurements per minute from right after the chitin treatment.

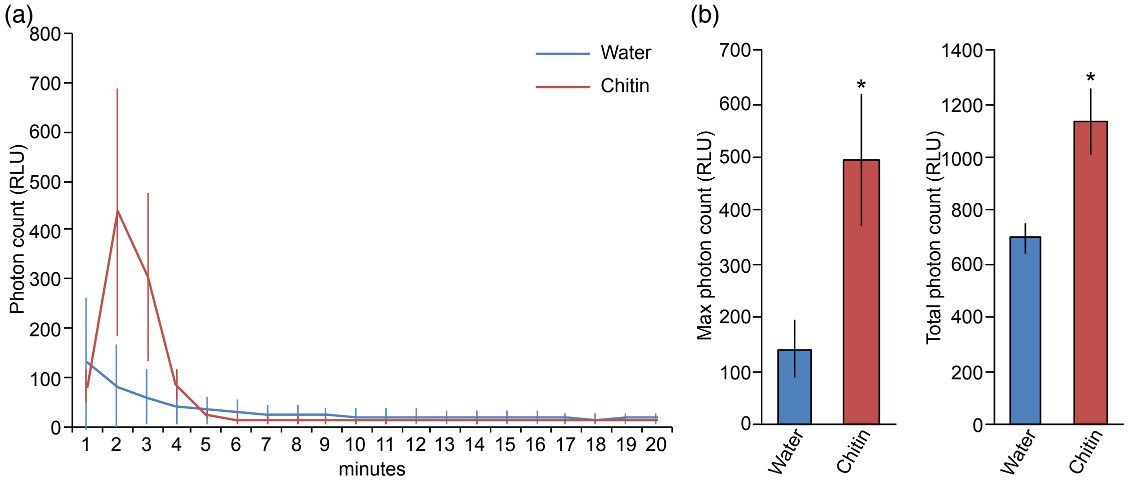

**Fig. S5** The two predicted glycosylation sites in SRN2 are not involved in the potentiation of chitin-triggered ROS bursts

(a) Schematics of SRN2$^{N101Q/N143Q}$-HA protein. SRN2 101st and 143rd asparagine (N) residues in the ribonuclease domain are predicted as glycosylation sites. (b) Expression of both SRN2-HA and SRN2$^{N101Q/N143Q}$-HA proteins was confirmed by immunoblot analysis. The upper and lower band of SRN2-HA correspond to two sites and single sites glycosylated SRN2-HA protein, respectively. The single band in the SRN2$^{N101Q/N143Q}$-HA lane represents the size of the non-glycosylated SRN2-HA protein. (c) The SRN2$^{H130A}$-HA protein is glycosylated. The deglycosylation enzyme treatment of the SRN2$^{H130A}$-HA protein expressed in *N. benthamiana* reduces the intensity of  the upper two bands and generates a lower band which is the same size as that of SRN2$^{N101Q/N143Q}$-HA. (d, e) Potentiation chitin-triggered ROS burst by SRN2 in *C. sativus* was not compromised by mutations in glycosylation sites. Experiments were performed in the same way as described in Fig. 4a. (f) SRN2$^{N101Q/N143Q}$-HA expression in *C. sativus* significantly increased chitin-triggered ROS burst. ** indicates $p<0.01$. (g) SRN2$^{N101Q/N143Q}$-HA expression in *C. sativus* showed no visible phenotype. Photographs were taken at 6 dpi. (b, c) Anti-HA antibody was used. CBB-stained RBCL proteins were used as loading controls.

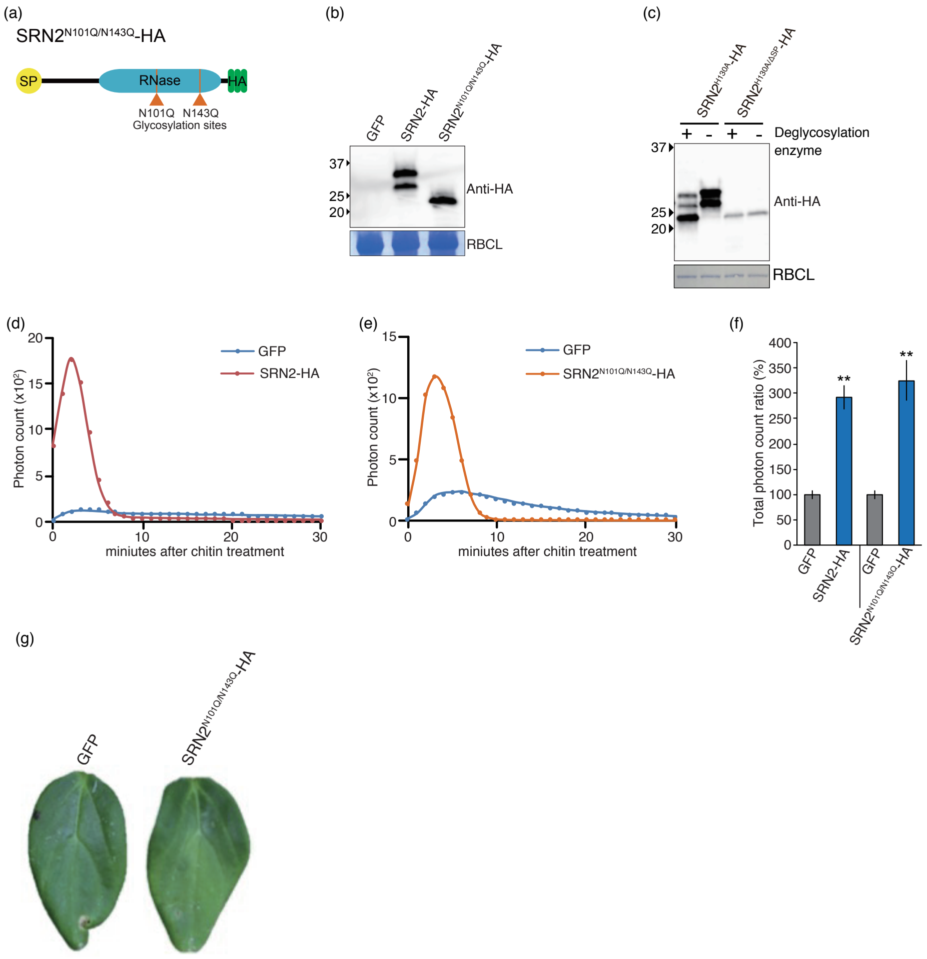

**Fig. S6** Assessment of PTI marker gene candidates in *C. sativus*

Accumulation of *C. sativus* PTI marker gene candidates' transcripts was quantified by RT-qPCR after chitin treatment. *Arabidopsis* PTI marker gene homologs were predicted by BLAST. *C. sativus* genes, Csa3G850580.1 and Csa6G454460.1 were strongly induced 30 minutes after chitin treatment and designated as *CsFRK1* and *CsNHL10*, respectively. *CsCYC* was used as an endogenous control. Data represent mean and ±SE (n=3). Two technical replicates are used. Primers used are listed in Supplementary information Table S2.

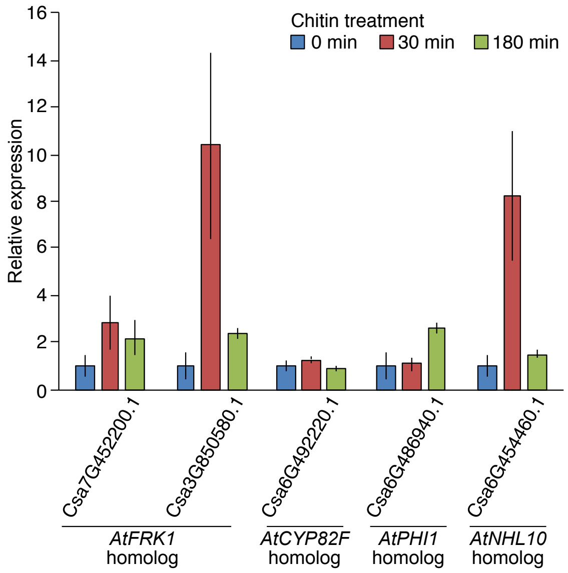

**Fig. S7** SRN1 does not cleave dsRNAs

(a) SRN1$^{H63A/\Delta SP}$ cleaves AG10 even without metal ions. In vitro RNase assay was performed in RNase buffer with EDTA. (b) SRN1$^{H63A/\Delta SP}$ generates laddered intermediated fragments of AG10. The molar ratios of SRN1 $^{H63A/\Delta SP}$ to 0.05 pmol AG10 was depicted on the top of image. (c) SRN1$^{H63A/\Delta SP}$ cleaves ssRNA4 but does not dsRNA4.

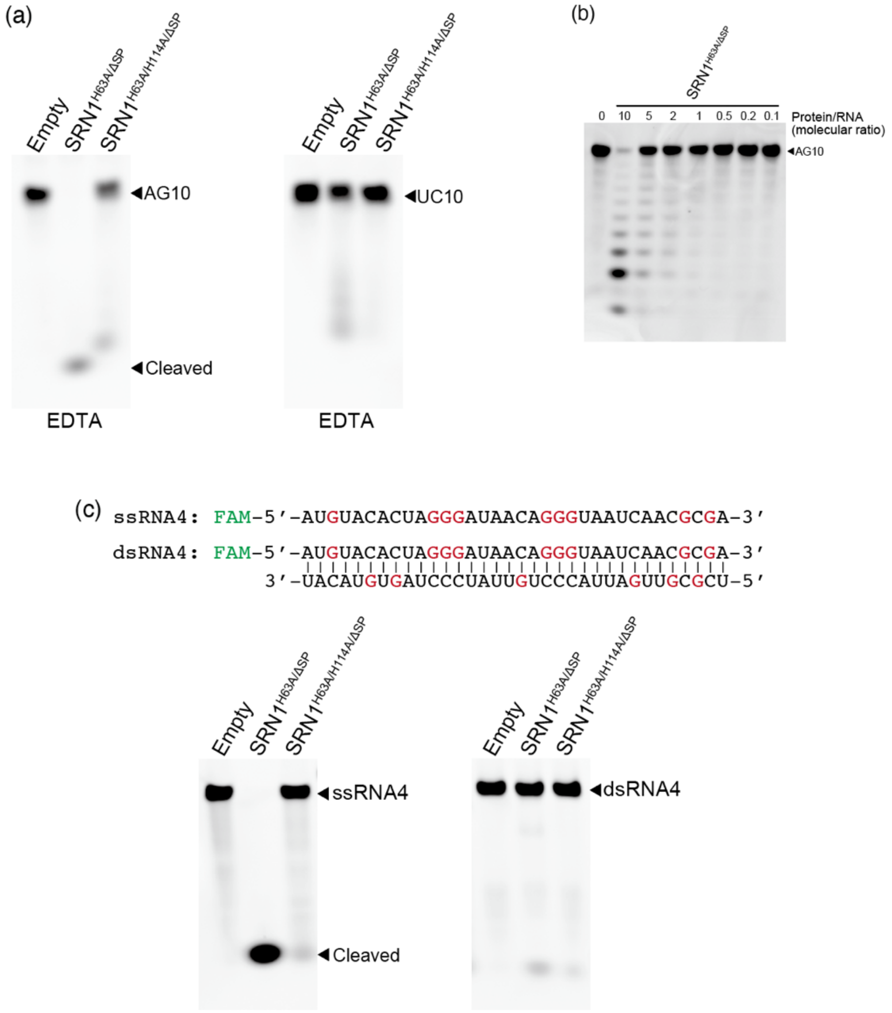

**Fig. S8** Phenotype of *N. benthamiana* expressing SRN2 mutant proteins
(a)SRN2-HA, SRN2$^{\Delta SP}$-HA, and SRN2$^{H130A}$-HA were expressed using an *Agrobacterium* mediated transient gene expression system. The photograph was taken at 5 dpi. (b) Immunoblot analysis of proteins isolated from *N. benthamiana*. Anti-HA antibody was used for detection of HA tagged proteins. Total proteins were extracted at 2 dpi. CBB-stained RBCL proteins were used as loading controls.

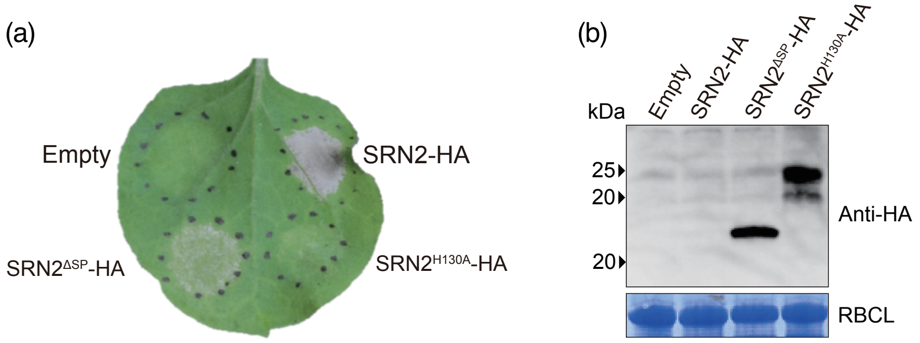

### **Table S1** List of *Colletotorichum orbiculare* conserved effector candidates

| # | *C. orbiculare* effector candidates: Name | Locus tag | Previous locus tag* | Annotation in NCBI | Homolog of *C. higginsianum*: Locus tag | Description |
|---|---|---|---|---|---|---|
| 1 | | Cob_v000569 | Cob_03109 | hypothetical protein | CH63R_01904 | |
| 2 | | Cob_v001255 | Cob_03592 | hypothetical protein | CH63R_11993 | |
| 3 | | Cob_v001256 | Cob_03593 | hypothetical protein | CH63R_11994 | |
| 4 | | Cob_v002378 | Cob_06769 | hypothetical protein | CH63R_09513 | |
| 5 | | Cob_v002526 | Cob_06927 | hypothetical protein | CH63R_11224 | |
| 6 | | Cob_v003377 | Cob_04472 | hypothetical protein | CH63R_10618 | |
| 7 | | Cob_v003509 | Cob_04605 | hypothetical protein | CH63R_03439 | Previously described as ChEC42** |
| 8 | | Cob_v005354 | Cob_06008 | hypothetical protein | CH63R_01410 | Previously described as ChEC20** |
| 9 | | Cob_v005447 | Cob_07794 | hypothetical protein | CH63R_13244 | |
| 10 | | Cob_v006915 | Cob_06145 | hypothetical protein | CH63R_06441 | |
| 11 | | Cob_v007014 | Cob_02253 | hypothetical protein | CH63R_05005 | |
| 12 | | Cob_v008050 | Cob_05125 | hypothetical protein | CH63R_01621 | |
| 13 | | Cob_v008419 | Cob_05507 | Carbonic anhydrase | CH63R_01803 | |
| 14 | | Cob_v009387 | Cob_08542 | hypothetical protein | CH63R_10425 | Previously described as ChEC51** |
| 15 | *SRN1* | Cob_v010174 | Cob_00225 | Guanyl-specific ribonuclease F1 | CH63R_08290 | |
| 16 | | Cob_v010422 | Cob_00825 | Tyrosinase ustQ | CH63R_05546 | |
| 17 | | Cob_v010630 | Cob_05661 | hypothetical protein | CH63R_11980 | Previously described as ChEC65** and ChCEC2-2*** |
| 18 | | Cob_v011474 | Cob_00326 | hypothetical protein | CH63R_05348 | |
| 19 | | Cob_v011718 | Cob_02689 | hypothetical protein | CH63R_08281 | |
| 20 | | Cob_v011797 | Cob_01861 | Polysaccharide monooxygenase Cel61a | CH63R_12153 | |
| 21 | | Cob_v012302 | Cob_06329 | hypothetical protein | CH63R_13188 | Previously discribed as ChEC6** |

*Gan et al., 2013, New Phytol., 197(4):1236
**Dallery et al, 2017, BMC Genomics, 18:667
***Tsushima et al., 2021, Front. in Microbiol., 12:2535

### **Table S2** List of 32 fungal genomes

| Species | Reference | Prefix | Phylum | Subphylum | Class |
|---|---|---|---|---|---|
| *Arthrobotrys oligospora* ATCC 24927 | Yang et al., 2011 | ART | Ascomycota | Pezizomycotina | Orbilomycetes |
| *Aspergillus nidulans* | Arnaud et al., 2012 | ASP | Ascomycota | Pezizomycotina | Eurotiomycetes |
| *Blumeria graminis* f. sp. *hordei* DH14 | Spanu et al., 2010 | BLU | Ascomycota | Pezizomycotina | Leotiomycetes |
| *Botrytis cinerea* | Amselem et al., 2011 | BOT | Ascomycota | Pezizomycotina | Leotiomycetes |
| *Chaetomium globosum* | Berka et al., 2011 | CHA | Ascomycota | Pezizomycotina | Sordariomycetes |
| *Cladonia grayi* Cgr/DA2myc/ss | JGI, Duke University | CLA | Ascomycota | Pezizomycotina | Lecaronomycetes |
| *Colletotrichum chlorophyti* | Gan et al., 2017 | CCH | Ascomycota | Pezizomycotina | Sordariomycetes |
| *Colletotrichum fioriniae* PJ7 | Baroncelli et al., 2014 | CFI | Ascomycota | Pezizomycotina | Sordariomycetes |
| *Colletotrichum fructicola* Nara gc5 | Gan et al., 2013 | CFR | Ascomycota | Pezizomycotina | Sordariomycetes |
| *Colletotrichum graminicola* | O'Connell et al., 2012 | CGR | Ascomycota | Pezizomycotina | Sordariomycetes |
| *Colletotrichum higginsianum* IMI 349063 | Zampounis et al., 2016 | CHI | Ascomycota | Pezizomycotina | Sordariomycetes |
| *Colletotrichum incanum* | Gan et al., 2016 | CIN | Ascomycota | Pezizomycotina | Sordariomycetes |
| *Colletotrichum orbiculare* MAFF 240422 104-T | Gan et al., 2013 | COR | Ascomycota | Pezizomycotina | Sordariomycetes |
| *Cryptococcus neoformans* JEC21 (PRJNA10698) | Loftus et al., 2005 | CRY | Basidiomycota | Agaricomycotina | Tremellomycetes |
| *Eutypa lata* UCR-EL1 | Blanco-Ulate et al., 2013 | EUT | Ascomycota | Pezizomycotina | Sordariomycetes |
| *Fusarium graminearum* PH-1 | Cuomo et al., 2007 | FUS | Ascomycota | Pezizomycotina | Sordariomycetes |
| *Leptosphaeria maculans* | Rouxel et al., 2011 | LEP | Ascomycota | Pezizomycotina | Dothideomycetes |
| *Magnaporthe oryzae* MG8 | Dean et al., 2005 | MOR | Ascomycota | Pezizomycotina | Sordariomycetes |
| *Metarhizium robertsii* ARSEF 23 | Hu et al., 2014 | MET | Ascomycota | Pezizomycotina | Sordariomycetes |
| *Nectria haematococca* MPV1, strain 77-13-4 | Coleman et al., 2009 | NEC | Ascomycota | Pezizomycotina | Sordariomycetes |
| *Neurospora crassa* OR74A | Galagan et al., 2003 | NEU | Ascomycota | Pezizomycotina | Sordariomycetes |
| *Podospora anserina* S Mat | Espagne et al., 2008 | POD | Ascomycota | Pezizomycotina | Sordariomycetes |
| *Puccinia graminis* f. sp. *tritici* | Duplessis et al., 2011 | PUC | Basidiomycota | Pucciniomycotina | Pucciniomycetes |
| *Rhizophagus irregularis* DAOM 181602 | Tisserant et al., 2013 | RHI | Glomeromycota | Glomeromycotina | Glomeromycetes |
| *Saccharomyces cerevisiae* S288C | Goffeau et al., 1996 | SAC | Ascomycota | Saccharomycotina | Saccharomycetes |
| *Taphrina deformans* | Cissé et al., 2013 | TAP | Ascomycota | Taphrinamycotina | Taphrinamycetes |
| *Trichoderma virens* Gv29-8 | Kubicek et al., 2011 | TRI | Ascomycota | Pezizomycotina | Sordariomycetes |
| *Tuber melanosporum* Mel28 | Martin et al., 2010 | TUB | Ascomycota | Pezizomycotina | Tuber melanosporum |
| *Ustilago maydis* | Kämper et al., 2006 | UST | Basidiomycota | Ustilaginomycotina | Ustilaginomycetes |
| *Verticillium dahliae* | Klosterman et al., 2011 | VER | Ascomycota | Pezizomycotina | Sordariomycetes |
| *Xylona heveae* TC161 | Gazis et al., 2016 | HYL | Ascomycota | Pezizomycotina | Xylonomycetes |
| *Zymoseptoria tritici* | Goodwin et al., 2011 | ZYM | Ascomycota | Pezizomycotina | Dothideomycetes |

### **Table S3** Plasmid list

| Name | Description | Reference |
| --- | --- | --- |
| pEAQ-HT-GFP | GFP expression in plants | (Sainsbury et al., 2009) |
| PBBR-gabT | For enhancing the T-DNA insertion from Agrobacteria to host plants | (Nonaka et al., 2017) |
| pENTR4-Cob_00225.1w/oSC | For the construction of pEAQ-Cob_00225.1 | This report |
| pENTR4-Cob_00565.1w/oSC | For the construction of pEAQ-Cob_00565.1 | This report |
| pENTR4-Cob_09372.1w/oSC | For the construction of pEAQ-Cob_09372.1 | This report |
| pENTR4-Cob_09372.2w/oSC | For the construction of pEAQ-Cob_09372.2 | This report |
| pENTR4-Cob_02744.1w/oSC | For the construction of pEAQ-Cob_02744.1 | This report |
| pENTR4-Cob_00565.1H130A | For the construction of pEAQ-Cob_00565.1H130A | This report |
| pENTR4-Cob_00565.1H75A/H130A | For the construction of pEAQ-Cob_00565.1H75A/H130A | This report |
| pENTR4-Cob_00565.1ΔSP | For the construction of pEAQ-Cob_00565.1SP | This report |
| pGWB2_Cob_00225 | *C. orbiculare* SRN1 expression in plants | This report |
| pGWB2_Cob_00565 | *C. orbiculare* SRN2 expression in plants | This report |
| pGWB2_Cob_09372.1 | *C. orbiculare* SRN3.1 expression in plants | This report |
| pGWB2_Cob_09372.2 | *C. orbiculare* SRN3.2 expression in plants | This report |
| pGWB2_Cob02744 | *C. orbiculare* SRN4 expression in plants | This report |
| pGWB2-GFP | GFP expression in plants | This report |
| pEAQ-Cob_00225.1 | For SRN1 expression in plants and for the construction of pNK046 | This report |
| pEAQ-Cob_00565.1 | For SRN2 expression in plants and for the construction of pNK047 | This report |
| pEAQ-Cob_09372.1 | For SRN3.1 expression in plants and for the construction of pNK048 | This report |
| pEAQ-Cob_09372.2 | For SRN3.2 expression in plants and for the construction of pNK049 | This report |
| pEAQ-Cob_02744.1 | For SRN4 expression in plants and for the construction of pNK052 | This report |
| pEAQ-Cob_00565.1H130A | For SRN2H130A expression in plants and for the construction of pNK040 | This report |
| pEAQ-Cob_00565.1H75A/H130A | For SRN2H75A/H130A expression in plants and for the construction of pNK093 | This report |
| pEAQ-Cob_00565.1ΔSP | For SRN2ΔSP expression in plants and for the construction of pNK060 | This report |
| pNK046_pEAQ-Cob_00225-3xHA | SRN1-HA expression in plants | This report |
| pNK047_pEAQ-Cob_00565.1-3xHA | SRN2-HA expression in plants | This report |
| pNK048_pEAQ-Cob_09372.1-3xHA | SRN3.1-HA expression in plants | This report |
| pNK049_pEAQ-Cob_09372.2-3xHA | SRN3.2-HA expression in plants | This report |
| pNK052_pEAQ-Cob_02744-3xHA | SRN4-HA expression in plants | This report |
| pNK130_pEAQ-Cob_00225.1H114A-3xHA | SRN1H114A-HA expression in plants | This report |
| pNK134_pEAQ-Cob_00225.1H63A/H114A-3xHA | SRN1H63A/H114A expression in plants | This report |
| pNK135_pEAQ-Cob_00225.1ΔSP--3xHA | SRN1ΔSP-HA expression in plants | This report |
| pNK040_pEAQ-Cob_00565.1H130A-3xHA | SRN2H130A-HA expression in plants | This report |
| pNK093_pEAQ-Cob_00565.1H75A/H130A-3xHA | SRN2H75A/H130A-HA expression in plants | This report |
| pNK060_pEAQ-Cob_00565.1ΔSP-3xHA | SRN2ΔSP-HA expression in plants | This report |
| pNK042_pEAQ-Cob_00565.1H130A/ΔSP-3xHA | SRN2H130A/ΔSP-HA expression in plants | This report |
| pNK102_pEAQ-Cob_00565.1N101Q/N143Q-3xHA | SRN2N101Q/N143Q-HA expression in plants | This report |
| pNK106_pENTR4-Ptef-Cob_00565_GntR(NPTII) | For SRN2 expression in *C. orbiculare* | This report |
| pNK109_pENTR4-Ptef-Cob_00565H75A/H130A_GntR(NPTII) | For SRN2H75A/H130A expression in *C. orbiculare* | This report |
| pNK144_pPICZA+a-factor_dSPCob_00225H63A-Pichia_myc-6xHis | For a-factor-SRN1H63A/ΔSP-myc-6xHis expression in Pichia pastoris | This report |
| pNK146_pPICZA+a-factor_dSPCob_00225H63AH114A-Pichia_myc-6xHis | For a-factor-SRN1H63A/H114A/ΔSP-myc-6xHis expression Pichia pastoris | This report |
| pKOSRN1 | For *C. orbiculare SRN1* (Cob_00225) knock-out | This report |
| pKOSRN2 | For *C. orbiculare SRN2* (Cob_00565) knock-out | This report |

### **Table S4** Oligonucleotide list

| Name | 5'-Sequence-3' | Description | Reference |
|---|---|---|---|
| 8436qF_ref | CTGGAACTGGGACTTGAAGC | *C. orbiculare* ribosomal protein L5 gene (Cob_v012718)quantification by RT-qPCR | (Gan *et al*., 2013) |
| 8436qR_ref | GCCGACACCCTGAAGAACTA | *C. orbiculare* ribosomal protein L5 gene (Cob_v012718)quantification by RT-qPCR | (Gan *et al*., 2013) |
| qCob_00225_3F | TCGTCTCTCTCTTCGCCATCGTG | SRN1 transcripts quantification by RT-qPCR | This report |
| qCob_00225_3R | AGGGCCCGGTGACATCAAAGTC | SRN1 transcripts quantification by RT-qPCR | This report |
| qCob_00565_2F | AGGGCCTCATCTTTGCCTCCACC | SRN2 transcripts quantification by RT-qPCR | This report |
| qCob_00565_2R | GCCCTGCCTGAGTAGTTTCCGGAC | SRN2 transcripts quantification by RT-qPCR | This report |
| qCob_09372.1_2F | CAATTCTGAGGGGTTCGAATTCCACGGTG | SRN3.1 transcripts quantification by RT-qPCR | This report |
| qCob_09372.1_2R | AGCTGTGATAATCCGCGCAAGACGC | SRN3.1 transcripts quantification by RT-qPCR | This report |
| qCob_09372.2_2F | TTCCGCAATTCTGAGGGGTTCGAATTCC | SRN3.2 transcripts quantification by RT-qPCR | This report |
| qCob_09372.2_2R | TAGATGCCCATCGTCTTCATCGGAAACTCG | SRN3.2 transcripts quantification by RT-qPCR | This report |
| qCob_02744_2F | TCTCCACGGAGAAGTTCGTCGTCAACG | SRN4 transcripts quantification by RT-qPCR | This report |
| qCob_02744_2R | ACTCTTGGAGGGGGTTCTTCTTGTCGTC | SRN4 transcripts quantification by RT-qPCR | This report |
| qCsCYC_F | GCTGGACCTGGAACCAACGGA | *C. sativus* reference gene *CYC* quantification by RT-qPCR | (Liang *et al*., 2018) |
| qCsCYC_R | TCTAAGAGAGCTGGCCACAAT | *C. sativus* reference gene *CYC* quantification by RT-qPCR | (Liang *et al*., 2018) |
| qCsEF1a_F1 | ACTGGTGGTTTTGAGGCTGGT | *C. sativus EF-1a* quantification by RT-qPCR | (Liang *et al*., 2018) |
| qCsEF1a_R1 | CTTGGAGTATTTGGGTGTGGT | *C. sativus EF-1a* quantification by RT-qPCR | (Liang *et al*., 2018) |
| qCsa7G452200.1_F1 | CAACATCATCATTCCAATTGTAGC | *C. sativus FRK1* homolog 1st quantification by RT-qPCR | This report |
| qCsa7G452200.1_R1 | TGCAACAATTGTGACAACAACTA | *C. sativus FRK1* homolog 1st quantification by RT-qPCR | This report |
| qCsa3G850580.1_F1 | ACGGTGGCATAATGAATCG | *C. sativus FRK1* homolog 2nd quantification by RT-qPCR | This report |
| qCsa3G850580.1_R1 | GGTGCTGTAAAACACCAATTCC | *C. sativus FRK1* homolog 2nd quantification by RT-qPCR | This report |
| qCsa6G492220.1_F2 | GAGCTGATTAAACAGATTGTGTCG | *C. sativus CYP82F2* 2nd homolog quantification by RT-qPCR | This report |
| qCsa6G492220.1_R2 | ACCCAATTCATCATCGGAAT | *C. sativus CYP82F2* 2nd homolog quantification by RT-qPCR | This report |
| qCsa6G486940.1_F1 | AGGTCATGAAACTGGCGAGA | *C. sativus PHI1* homolog quantification by RT-qPCR | This report |
| qCsa6G486940.1_R1 | ATTTGATTCGCTGTGATTCTGA | *C. sativus PHI1* homolog quantification by RT-qPCR | This report |
| qCsa6G454460.1_F4 | GGCTCATCTTCCGTCCAA | *C. sativus NHL10*, 3rd homolog quantification by RT-qPCR | This report |
| qCsa6G454460.1_R4 | GGTGAAATTGAACTGCGTGA | *C. sativus NHL10*, 3rd homolog quantification by RT-qPCR | This report |
| IF-Cob_00225+pENTR4_F | GCAGGCTCCACCATGCAGTTCTCCGTCGTCCTC | Construction | This report |
| IF-Cob_00225(w/oSC)+pENTR4_R | AAGCTGGGTCTAGATAGAGGTACCAGAGCATC | Construction | This report |
| IF-Cob_00565+pENTR4_F | GCAGGCTCCACCATGATCAGCCTCAAATCGACGC | Construction | This report |
| IF-Cob_00565(w/oSC)+pENTR4_R | AAGCTGGGTCTAGATCGCGATGAAGGCCAT | Construction | This report |
| IF-Cob_09372+pENTR4_F | GCAGGCTCCACCATGTTCTTAGGGGTAGCCATCA | Construction | This report |
| IF-Cob_09372(w/oSC)_R | AAGCTGGGTCTAGATGCTGTGATAATCCGCGCAA | Construction | This report |
| IF-pETNR4-Cob_09372.2 woSC_R | AAGCTGGGTCTAGATGGGGTTCCCGCCAGTGTAG | Construction | This report |
| IF-Cob_02744+pENTR4_F | GCAGGCTCCACCATGCACTCCACCAAGATCCTC | Construction | This report |
| IF-Cob_02744(w/oSC)_R | AAGCTGGGTCTAGATGTCGCACTTCTGGAAGTAGT | Construction | This report |
| Cob_00565H130A_R | GCGCCGGTGGCGGTCATTGCGCCGACGTAG | Construction | This report |
| Cob_00565H130A_F | GCAATGACCGCCACCGGCGCTGACCAGCGC | Construction | This report |
| Cob_00565.1_H58A_mut.R | AAGCGAGCCGGGTACTGGCTGGT | Construction | This report |
| Cob_00565.1_H58A_mut.F | AGCCAGTACCCGGCTCGCTTCA | Construction | This report |
| IF-noSP-Cob_00565_F | GCAGGCTCCACCATGTCGCCACTGCCCGAACAGACCCT | Construction | This report |
| IF-GFP+pENTR4_F | GCAGGCTCCACCATGGTGAGCAAGGGCGAGGAG | Construction | This report |
| IF-GFP+pENTR4_R | AAGCTGGGTCTAGATTTACTTGTACAGCTCGTCC | Construction | This report |
| HF-pEAQ-HT_F | CTCGAGGCCTTTAACTCTGGTTTCATTAAATTTTC | Construction | This report |
| IF-pEAQHTCob_00565_R | TCCGCCGGATCCGCCCGCGATGAAGGCCATGAAAGAC | Construction | This report |
| IF-Cter-GGSGG-HA_F | GGCGGATCCGGCGGATACCCATACGATGTTCCTGACTATGC | Construction | This report |
| HF-GGSGG-3xHA_R | TGAAACCAGAGTTAAAGGCCTCGAGTTAAGCGTAATCTGGAACGTCATATGG | Construction | This report |
| HF-Cob_00225_R | GGGTATCCGCCGGATCCGCCAGAGGTACCAGAGCATC | Construction | This report |
| HF-Cob_09372.1_R | GGGTATCCGCCGGATCCGCCGCTGTGATAATCCGCGCAA | Construction | This report |
| HF-Cob_09372.2_R | GGGTATCCGCCGGATCCGCCGGGGTTCCCGCCAGTGTAG | Construction | This report |
| HF-Cob_02744_R | GGGTATCCGCCGGATCCGCCGTCGCACTTCTGGAAGTAGT | Construction | This report |
| HF-pNK102BB_F2 | GAGACGCGGAGGGGCGGCA | Construction | This report |
| HF-pNK102BB_R2 | TCCGGACGGTATGATGGGAAACTC | Construction | This report |
| HF-pNK102IN_F2 | ATCATACCGTCCGGACAGTACTCAGGCAGG | Construction | This report |
| HF-pNK102IN_R2 | GCCCCTCCGCGTCTCCTGGCACGTCACGAAGCC | Construction | This report |
| HF-pNK073IN_F | AACTCTTCCTGTTAGTTAGTGACCCTTTGGCTCGCTTAG | Construction | This report |
| HF-pNK073IN_R | GATAGCTGTCGCTGTCAACTCCCCCTATCCCATAAATGC | Construction | This report |
| HF-pNK072_F | AGTTGACAGCGACAGCTATCAGTTG | Construction | This report |
| HF-pNK072_R | ACTAACTAACAGGAAGAGTTTGTAGAAACGC | Construction | This report |
| HF-pNK106forGntR_F | AATCCTTCTTCCATCCAGCTGCAGCTCT | Construction | This report |
| HF-pNK106forGntR_R | CAACTGGCATTTCACCAGCCCCTGTTCTCG | Construction | This report |
| HF-GntRpNK106_F | GGCTGGTGAAATGCCAGTTGTTCCCAGTGATC | Construction | This report |
| HF-GntRpNK106_R | AGCTGGATGGAAGAAGGATTACCTCTAAACAAGTGTACC | Construction | This report |
| HF-pNK1062nd_F | TTCATCGCGTGAGGATCCCCCGGGCTGCAGGA | Construction | This report |
| HF-pNK07X-BB_R | TGTGTCTGCCTCTAGAGTGATGTATGGAAGATGGAGTG | Construction | This report |
| HF-pNK073+SRN2_F | CTCTAGAGGCAGACACAATGATCAGCCTCAAATCGACGC | Construction | This report |
| HF-mut-Cob_00225H114A_F | GCAGATCACGGCCACTGGAGCGAGCGGTAAC | Construction | This report |
| HF-mut-Cob_00225H114A_R | CTCCAGTGGCCGTGATCTGCCCAGCCTGG | Construction | This report |
| HF-pNK133H63A_F | CGTACCCCGCTCGCTACAACAACTACGAGGGC | Construction | This report |
| HF-pNK133H63A_R | TTGTAGCGAGCGGGGTACGTCGACGTACCG | Construction | This report |
| HF-pNK135_F | CGCGACCGGTATGGTCCCCACGGAACTGCA | Construction | This report |
| HF-pNK135_R | TGGGGACCATACCGGTCGCGAATTTGGGCA | Construction | This report |
| CoSRN1_UPfwd | GACTGCAGCGGATTATTGCTAGGATTTGCCGA | Construction | This report |
| CoSRN1_UPrev | AACTGCAGTTTGGAAGCAGGCGAGGATGG | Construction | This report |
| CoSRN1_DWfwd | CGGGGTACCATGAGCTCGAGTGCGAACGGG | Construction | This report |
| CoSRN1_DWrev | CGCGGGTACCCGGCAACGTGGCATCGTGGCA | Construction | This report |
| CoSRN2_UPfwd | TATAGGGCCCGACCCGTGGGAAGCTCGAGACGAC | Construction | This report |
| CoSRN2_UPrev | TATAGGGCCCGGTGATGGGCGAGTGCGGAGGCGA | Construction | This report |
| CoSRN2_DWfwd | GCGCGCGGCCGCCAAGACTTGTTTTCGATCTGGTTG | Construction | This report |
| CoSRN2_DWrev | GCGCGCGGCCGCGGCTGATCTCTTGGCTCTTCTTCG | Construction | This report |
| HF-pPCIZBB_F | GAACAAAAACTCATCTCAGAAGAGGATCTGAATAG | Construction | This report |
| HF-pPCIZBB_R | CGTTTCGAATAATTAGTTGTTTTTTGATCTTCTCAAG | Construction | This report |
| HF-alpha_F | ACAACTAATTATTCGAAACGATGAGATTTCCTTCAATTTTTACTGCTGTTTTATTCG | Construction | This report |
| HF-alpha_R | AGCTTCAGCCTCTCTTTTCTCGAGA | Construction | This report |
| HF-pNK144IN_F | GGCTGAAGCTATGGTACCAACGGAATTGCAGAAG | Construction | This report |
| HF-pNK144IN_R | CGGCGCTATTCAGATCCTCTTCTGAGATGAGTTTTTGTTCGCCTCCGCTACCACCACTGG | Construction | This report |
| HF-pNK146_F | ATAACCGCTACAGGTGCTTCTGGGAACAACTTC | Construction | This report |
| HF-pNK146_R | GAAGCACCTGTAGCGGTTATCTGTCCTGCTTGTCCACA | Construction | This report |
| Linker oligonucleotide | (Phos)ATCGTATCGTAGATCGGAAGAGCACACGTCTGAA(ddC) | Linker ligation assay. (Phos) and (ddC) indicates a 5'-end phosphate and a 3' terminal 2′, 3′-dideoxycytidine, respectively | This report |

**Table S5** Fungal strain list

| Name | Genotype |
|---|---|
| *C. orbiculare* 104-T | Wild-type (MAFF240422) |
| *srn1*#21 | *srn1::HPH* |
| *srn1*#25 | *srn1::HPH* |
| *srn2*#8 | *srn2::SUR* |
| *srn2*#11 | *srn2::SUR* |
| *srn1 srn2*#1 | *srn1::HPH srn2::SUR* |
| *srn1 srn2*#2 | *srn1::HPH srn2::SUR* |
| *Tef:SRN2 in srn12*#1 | *Tef:SRN2 srn1::HPH srn2::SUR* |
| *Tef:SRN2 in srn12*#2 | *Tef:SRN2 srn1::HPH srn2::SUR* |
| *Tef:SRN2H75A/H130A in srn12*#1 | *Tef:SRN2H75A/H130A srn1::HPH srn2::SUR* |
| *Tef:SRN2H75A/H130A in srn12*#2 | *Tef:SRN2H75A/H130A srn1::HPT srn2::SUR* |

**Table S6** Accession numbers of *SRN* genes

| Gene name | Locus tag* | Previous locus tag** | GenBank | Annotation in NCBI |
|---|---|---|---|---|
| *SRN1* | Cob_v010174 | Cob_00225 | TDZ16998.1 | Guanyl-specific ribonuclease F1 |
| *SRN2* | Cob_v012824 | Cob_00565 | TDZ14290.1 | Guanyl-specific ribonuclease F1 |
| *SRN3* | Cob_v000102 | Cob_09372 | TDZ26080.1 | Extracellular guanyl-specific ribonuclease |
| *SRN4* | Cob_v009749 | Cob_02744 | TDZ17187.1 | Ribonuclease clavin |

**C. orbiculare* MAFF 240422 104-T (NCBI accession: PRJNA171217), reffered in Gan et al., 2019, *Mol. Plant-Microbe Inter.*, 32(9)
**Reffered in Gan et al., 2013, *New Phytol.*, 197(4):1236

**Table S7** Number of *SRN* genes in *Colletotrichum* species

| Species complex | Species | *SRN1/3* | *SRN2* | *SRN4* | others | total |
|---|---|---|---|---|---|---|
| acutatum | *C. acutatum* | 2 | 1 | 0 | 0 | 3 |
| | *C. nymphaeae* | 2 | 1 | 0 | 0 | 3 |
| | *C. fioriniae* | 2 | 1 | 0 | 0 | 3 |
| | *C. godetiae* | 2 | 1 | 0 | 0 | 3 |
| | *C. salicis* | 2 | 1 | 0 | 0 | 3 |
| | *C. simmondsii* | 2 | 1 | 0 | 0 | 3 |
| graminicola | *C. falcatum* | 2 | 1 | 0 | 0 | 3 |
| | *C. graminicola* | 2 | 1 | 0 | 0 | 3 |
| | *C. sublineola* | 2 | 1 | 0 | 0 | 3 |
| spaethianum | *C. incanum* INC03 | 1 | 1 | 0 | 1 | 3 |
| | *C. torieldiae* | 2 | 1 | 0 | 0 | 3 |
| destructivum | *C. higginsianum* CHjp | 1 | 1 | 0 | 0 | 2 |
| - | *C. chlorophyti* NTL11 | 1 | 1 | 0 | 0 | 2 |
| gloeosporioides | *C. aenigma* Cg56* | 2 | 2 | 1 | 0 | 8 |
| | *C. fructicola* 254* | 2 | 2 | 1 | 0 | 8 |
| | *C. siamense* CAD1* | 2 | 2 | 1 | 3 | 8 |
| | *C. tropicale** | 2 | 2 | 1 | 3 | 8 |
| orbiculare | *C. lindemuthianum* | 2 | 1 | 1 | 0 | 4 |
| | *C. malvarum* MAL* | 2 | 1 | 1 | 0 | 4 |
| | *C. orbiculare* 104T | 2 | 1 | 1 | 0 | 4 |
| | *C. spinosum* XAN* | 2 | 1 | 1 | 0 | 4 |
| | *C. triforii** | 2 | 1 | 1 | 0 | 4 |

*Unpublished genomes

**Method S1** Plasmids construction

All primers used are listed in Supporting Information Table S3. The genomic DNA of *C. orbiculare* 104-T and cDNA used for PCR was isolated and synthesized as previously described (Gan *et al.*, 2013).

**pENTR4-Cob_00225.1w/oSC, pENTR4-Cob_00565.1w/oSC, pENTR4-Cob_09372.1w/oSC, pENTR4-Cob_09372.2w/oSC, and pENTR4-Cob_02744.1w/oSC**

The Gateway pENTR4 Dual Selection Vector (Thermo Fisher Scientific) was linearized with NcoI and EcoRV, resulting in a pENTR4-NcoI-EcoRV fragment. PCR fragments (PCR-1 to -5) were amplified from cDNA synthesized with total RNA of *C. orbiculare* conidia using primer sets IF-Cob_00225+pENTR4_F plus IF-Cob_00225(w/oSC)+pENTR4_R, IF-Cob_00565+pENTR4_F plus IF-Cob_00565(w/oSC)+pENTR4_R, IF-Cob_09372+pENTR4_F plus IF-Cob_09372(w/oSC)_R, IF-Cob_09372+pENTR4_F plus IF-pETNR4-Cob_09372.2woSC_R, and IF-Cob_02744+pENTR4_F plus IF-Cob_02744(w/oSC)_R. pENTR4-NcoI-EcoRV and PCR-1 to -5 fragments were mixed and carried out In-Fusion cloning reaction using In-Fusion HD Cloning Kit (Takara), generating pENTR4-Cob_00225.1w/oSC, pENTR4-Cob_00565.1w/oSC, pENTR4-Cob_09372.1w/oSC, pENTR4-Cob_09372.2w/oSC, and pENTR4-Cob_02744.1w/oSC.

**pENTR4-Cob_00565.1$^{H130A}$**

PCR fragments (PCR-6 and -7) were amplified from pENTR4-Cob_00565.1w/oSC using primer sets IF-Cob_00565+pENTR4_F plus Cob_00565H130A_R and Cob_00565H130A_F plus IF-Cob_00565(w/oSC)+pENTR4_R. PCR-6 and PCR-7 were then mixed, and amplified with a primer set IF-Cob_00565+pENTR4_F plus IF-Cob_00565(w/oSC)+pENTR4_R, resulting in PCR-8. PCR-8 was mixed with pENTR4-NcoI-EcoRV and In-Fusion cloning was carried out using the In-Fusion HD Cloning Kit (Takara) according to the manufacturer's protocol, generating pENTR4-Cob_00565.1$^{H130A}$.

**pENTR4-Cob_00565.1$^{H75A/H130A}$**

PCR-9 and PCR-10 were amplified using primer sets IF-Cob_00565+pENTR4_F plus Cob_00565.1_H58A_mut.R and Cob_00565.1_H58A_mut.F plus IF-Cob_00565H130A(w/oSC)+pENTR4_R from pENTR4-Cob_00565.1w/oSC. The mixture of PCR-7 and -8 was amplified using primer set IF-Cob_00565+pENTR4_F plus IF-

Cob_00565(w/oSC)+pENTR4_R, resulting in PCR-10. An In-Fusion reaction was carried out using a mixture of PCR-10 and pENTR4-NcoI-EcoRV to obtain pENTR4-Cob_00565.1$^{H75A/H130A}$.

**pENTR4-Cob_00565.1$^{\Delta SP}$**

A PCR fragment (PCR-11) was amplified from pENTR4-Cob_00565.1w/oSC using primer set IF-noSP-Cob_00565_F plus IF-Cob_00565(w/oSC)+pENTR4_R. PCR-11 was then mixed with pENTR4-NcoI-EcoRV and In-Fusion cloning was carried out, generating pENTR4-Cob_00565.1$^{\Delta SP}$.

**pGWB2-Cob_00225, pGWB2-Cob_00565, pGWB2-Cob_09372.1, pGWB2-Cob_09372.2, and pGWB-Cob_02744**

pGWB2-Cob_00225, pGWB2-Cob_00565, pGWB2-Cob_09372.1, pGWB2-Cob_09372.2, pGWB-Cob_02744 were generated by an LR reaction mixing the destination vector pGWB2 (Nakagawa *et al.*, 2007) and entry vectors, pENTR4-Cob_00225.1w/oSC, pENTR4-Cob_00565.1w/oSC, pENTR4-Cob_09372.1w/oSC, pENTR4-Cob_09372.2w/oSC, and pENTR4-Cob_02744.1w/oSC, respectively.

**pGWB2-GFP**

A PCR fragment (PCR-12) was amplified from the pGWB5 plasmid (Nakagawa *et al.*, 2007) using primer set IF-GFP+pENTR4_F plus IF-GFP+pENTR4_R. PCR-12 and the pENTR4-NcoI-EcoRV fragment was assembled using the In-Fusion HD Cloning Kit (Takara), resulting in pENTR4-GFP. pGWB2-GFP was generated by the LR reaction mixing the destination vector pGWB2 (Nakagawa *et al.*, 2007) and an entry vector, pENTR4-GFP.

**pEAQ-Cob_00225.1, pEAQ-Cob_00565.1, pEAQ-Cob_09372.1, pEAQ-Cob_09372.2, pEAQ-Cob_02744.1, pEAQ-Cob_00565.1$^{H130A}$, pEAQ-Cob_00565.1$^{H75A/H130A}$, and pEAQ-Cob_00565.1$^{\Delta SP}$**

pEAQ-Cob_00225.1, pEAQ-Cob_00565.1, pEAQ-Cob_09372.1, pEAQ-Cob_09372.2, pEAQ-Cob_02744.1, pEAQ-Cob_00565.1$^{H130A}$, pEAQ-Cob_00565.1$^{H75A/H130A}$, and pEAQ-Cob_00565.1$^{\Delta SP}$ were generated by the LR reaction mixing a destination vector, pEAQ-HT-DEST1 (Sainsbury *et al.*, 2009), and each entry vector (pENTR4-Cob_00225.1 w/oSC, pENTR4-Cob_00565.1w/oSC, pENTR4-Cob_09372.1w/oSC, pENTR4-Cob_09372.2w/oSC, pENTR4-Cob_02744.1w/oSC, pENTR4-Cob_00565.1$^{H130A}$, pENTR4-Cob_00565.1$^{H75AH130A}$, and pENTR4 -Cob_00565.1$^{\Delta SP}$) using the Gateway LR Clonase II Enzyme Mix (Thermo Fisher Scientific), according to the manufacturer's protocol.

**pNK047, pNK040, pNK093, pNK060, and pNK042**

PCR-13 to -16 were amplified from pEAQ-Cob_00565.1, pEAQ-Cob_00565.1$^{H130A}$, pEAQ-Cob_00565.1$^{H75A/H130A}$, and pEAQ-Cob_00565.1$^{\Delta SP}$ using primer set IF-pEAQHT_F plus IF-pEAQHTCob_00565_R. A PCR fragment (PCR-17) encoding a linker peptide (GGSGG) and three HA tags derived from the human influenza hemagglutinin were amplified from pGWB14 (Nakagawa *et al.*, 2007) using primer set, IF-Cter-GGSGG-HA_F plus HF-GGSGG-3xHA_R. Each PCR13-16 product was mixed with PCR-17 and In-Fusion cloning was carried out, generating pNK047_pEAQ-Cob_00565.1-3xHA, pNK040_pEAQ-Cob_00565.1$^{H130A}$-3xHA, pNK093_pEAQ-Cob_00565.1$^{H75A/H130A}$-3xHA, and pNK060_pEAQ-Cob_00565.1$^{\Delta SP}$-3xHA. PCR-18 was amplified from pNK040 using primer set IF-noSP-Cob_00565_F plus HF-pENTR4BB-R. PCR-18 was then circularized by the In-Fusion cloning reaction, generating pNK042_pEAQ-Cob_00565.1$^{H130A/\Delta SP}$-3xHA.

**pNK046, pNK048, pNK049, and pNK052**

Four PCR fragments (PCR-19 to -22) were amplified from pEAQ-Cob_00225.1, pEAQ-Cob_09372.1, pEAQ-Cob_09372.2, pEAQ-Cob_02744.1 using primer sets HF-pEAQ-HT_F plus HF-Cob_00225_R, HF-pEAQ-HT_F plus HF-Cob_09372.1_R, HF-pEAQ-HT_F plus HF-Cob_09372.2_R, and HF-pEAQ-HT_F plus HF-Cob_02744_R, respectively. Each product of PCR-19 to -22 was mixed with PCR-17 and In-Fusion cloning was carried out, generating pNK046_pEAQ-Cob_00225-3xHA, pNK048_pEAQ-Cob_09372.1-3xHA, pNK049_pEAQ-Cob_09372.2-3xHA, pNK052_pEAQ-Cob_02744-3xHA.

**pNK102**

A PCR fragment (PCR-23) for the backbone of pNK102 was amplified from pNK047_pEAQ-Cob_00565.1-3xHA using primer set HF-pNK102BB_F2 plus HF-pNK102BB_R2. A PCR fragment (PCR-24) was amplified from pNK047_pEAQ-Cob_00565.1-3xHA using primer set HF-pNK102IN_F2 plus HF-pNK102IN_R2 to introduce the H101Q and H143Q mutations. PCR-23 and PCR-24 were then assembled using the NEBuilder HiFi DNA Assembly Master Mix (New England Biolabs) according to the manufacturer's protocol, resulting in pNK102_ pEAQ-Cob_00565.1$^{N101Q/N143Q}$-3xHA.

**pNK106 and pNK109**

A PCR fragment (PCR-25) containing the GFP expression cassette driven by the *Tef* promoter and *SCD1* terminator was amplified from TEF-GFP (Kumakura *et al.*, 2019) using primer set HF-pNK073IN_F plus HF-pNK073IN_R. A PCR fragment (PCR-25) for the backbone was amplified from pENTR4 using primer set HF-pNK072_F plus HF-pNK072_R. PCR-24 and PCR-25 were assembled using the NEBuilder HiFi DNA Assembly Master Mix, resulting in pETNR4-TefGFP. A PCR fragment (PCR-27) was amplified from pENTR4-TefGFP using primer set HF-pNK106forGntR_F plus HF-pNK106forGntR_R. The expression cassette of the Geneticin/G418 resistance gene, *NPTII*, was amplified from the pII99 plasmid (Namiki *et al.*, 2001) using primer set, HF-GntRpNK106_F plus HF-GntRpNK106_R, and named PCR-27. PCR-27 and PCR-28 were assembled using the NEBuilder HiFi DNA Assembly Master Mix (New England Biolabs), resulting in pENTR4-TefGFP-GntR. A PCR fragment (PCR-29) was amplified from pENTR4-TefGFP-GntR using primer set HF-pNK1062nd_F plus HF-pNK07X-BB_R. PCR fragments (PCR-30 and -31) were amplified using primer set HF-pNK073+SRN2_F plus HF-pNK073IN_R, from pNK047 and pNK093, respectively. PCR-32 and PCR-33 were assembled using the NEBuilder HiFi DNA Assembly Master Mix (New England Biolabs), resulting in pNK106_ pENTR4-Ptef-Cob_00565_GntR(NPTII). PCR-29 and PCR-31 were assembled using the NEBuilder HiFi DNA Assembly Master Mix, resulting in pNK109_pENTR4-Ptef-Cob_00565$^{H75A/H130A}$_GntR(NPTII).

**pNK130, pNK134, and pNK135**

PCR-31 was amplified using primer set HF-mut-Cob_00225H114A_F plus HF-mut-Cob_00225H114A_R, from pNK046. PCR-32 was then circularized by the NEBuilder HiFi DNA Assembly Master Mix, resulting in pNK130_pEAQ-Cob_00225.1$^{H114A}$-3xHA. PCR-33 was amplified using primer set HF-pNK133H63A_F plus HF-pNK133H63A_R, from pNK130. PCR-33 was then circularized by the NEBuilder HiFi DNA Assembly Master Mix, resulting in pNK134_pEAQ-Cob_00225.1H63A$^{H114A}$-3xHA. PCR-34 was amplified using primer set HF-pNK135_F plus HF-pNK135_R. PCR-34 was then circularized by the NEBuilder HiFi DNA Assembly Master Mix, resulting in pNK135_pEAQ-Cob_00225.1-3$^{\Delta SP}$-3xHA.

**pNK144 and pNK146**

PCR-35 was amplified using primer set HF-pPCIZBB_F plus HF-pPCIZBB_R, from pPICZ A (Thermo Fischer Scientific), a recombinant protein expression vector in *Pichia pastoris*. Nucleotide sequences of alpha-factor

(ATGAGATTTCCTTCAATTTTTACTGCTGTTTTATTCGCAGCATCCTCCGCATTAGC TGCTCCAGTCAACACTACAACAGAAGATGAAACGGCACAAATTCCGGCTGAAGC TGTCATCGGTTACTCAGATTTAGAAGGGGATTTCGATGTTGCTGTTTTGCCATTTT CCAACAGCACAAATAACGGGTTATTGTTTATAAATACTACTATTGCCAGCATTGC TGCTAAAGAAGAAGGGGTATCTCTCGAGAAAAGAGAGGCTGAAGCT), N-terminal secretion signal from *S. cerevisiae* and, codon optimized SRN1$^{\Delta SP/H63A}$ were synthesized by Eurofins genomics. PCR-36 and PCR-37 were amplified by primer sets, HF-alpha_F plus HF-alpha_R and HF-pNK144IN_F plus HF-pNK144IN_R from plasmid containing alpha-factor and SRN1$^{\Delta SP/H63A,}$ respectively. PCR-35, -36, and -37 were assembled using the NEBuilder HiFi DNA Assembly Master Mix, resulting in pNK144_ pPICZA+a-factor_dSPCob_00225H63A-Pichia_myc-6xHis. To introduce H114A mutation in pNK144, PCR-38 was amplified using a primer set, HF-pNK146_F plus HF-pNK146_R. PCR-38 was then circularized by the NEBuilder HiFi DNA Assembly Master Mix, resulting in pNK146_pPICZA+a-factor_dSPCob_00225H63A/H114A-Pichia_myc-6xHis.

**pKOSRN1 and pKOSRN2**

To construct the *CoSRN1* and *CoSRN2* replacement vectors, the 2 kb fragment of the 5' flanking region of *CoSRN1* was amplified by PCR using primer CoSRN1_UPfwd and primer CoSRN1_UPrev containing a PstI site. The amplified product was digested with PstI and introduced into the PstI site of pCB1636 harboring a hygromycin-resistance gene (*HPH*) to generate plasmid pKOSRN1_UP. Next, the 2 kb fragment of the 3' flanking region of *CoSRN1* was amplified using PCR with primer CoSRN1_DWfwd and primer CoSRN1_DWrev which contained KpnI. The amplified PCR product was digested with KpnI and introduced to the KpnI site of pKOSRN1_UP, resulting in pKOSRN1. The 2 kb fragment of the 5' flanking region of *CoSRN2* was amplified by PCR using primer CoSRN2_UPfwd and primer CoSRN2_UPrev containing an ApaI site. The ApaI-digested amplicon was ligated to the ApaI site of pCB1551 harboring the acetolactate synthase gene (*SUR*) (Sweigard *et al.*, 1997) to produce plasmid pKOSRN2_UP. The 2 kb fragment of the 3' flanking region of *CoSRN2* was amplified using PCR with primer CoSRN2_DWfwd and primer CoSRN2_DWrev which contained NotI. The amplified PCR product was digested with NotI and introduced to the NotI site of pKOSRN2_UP, resulting in pKOSRN2.

**Method S2** Fungal transformation

Fungal transformation was carried out using PEG-mediated protoplast transformation as described previously (Kubo *et al.*, 1991). Single knockout mutants, *srn1* and *srn2,* were generated by individually introducing pKOSRN1 and pKOSRN2 into *C. orbiculare*. Hygromycin B at 100 μg $ml^{-1}$ in PDA was used to select pKOSRN1 transformants. Sulfonylurea at 100 μg $ml^{-1}$ in a minimal medium (0.6 M glucose, 0.16 % yeast Nitrogen Base w/o Amino Acids, 0.2 % asparagine, 0.1 % ammonium nitrate, pH6.0) was used to select pKOSRN2 transformants. The *srn1 srn2* double knockout mutants were generated by transforming pKOSRN1 into the *srn2* strain. To establish *Tef:SRN2* and *Tef:SRN2*$^{H75A/H130A}$ in the *srn1 srn2* strains, pNK106_pENTR4-Ptef-Cob_00565_GntR and pNK109_pENTR4-Ptef-Cob_00565$^{H75A/H130A}$_GntR were used to transform *srn1 srn2#1* strains, respectively. G418 sulfate (Wako) at 200 μg $ml^{-1}$ was used for selection. Fungal strains used in this report are listed in Supporting Information Table S5.

**Method S3** Fungal inoculation

All the *C. orbiculare* strains were maintained on PDA medium in 90 × 15 mm plastic dishes (LMS Co., Ltd) at 25 ˚C in the dark. Three pieces of fungi grown on PDA were hollowed out using a plastic straw and transferred onto a new PDA dish and incubated at 25 ˚C for 6 d. For invasion assays, 5 μl of conidial solution at 1 × $10^5$ conidia $ml^{-1}$ was drop-inoculated onto detached *C. sativus* cotyledons at 6 days post germination (dpg). The leaves were then maintained in plastic dishes with paper towels moistened with water and incubated with high humidity at 24 ˚C under 10 h light:14 h dark conditions for 60 h. For fungal biomass measurements, conidial suspensions at 1 × $10^5$ conidia $ml^{-1}$ were sprayed onto detached *C. sativus* leaves at 9 dpg. Leaves were then incubated in the same way as were the drop-inoculated samples until sampling for RNA isolation.

**Method S4** Immunoblotting

For HA (from human influenza hemagglutinin)-tagged and GFP-fused protein isolation, leaf disks of *C. sativus* or *N. benthamiana* were ground and homogenized in buffer (150 mM NaCl, 1% (w $v^{-1}$) NP-40, 5 % glycerol, 50 mM Tris-HCl pH8.0, 5 mM DTT) with Protease Inhibitor Cocktail for plant cell and tissue extracts (P9599, Sigma-Aldrich) at 100 times dilution. To detect levels of MAPK phosphorylation, leaf disks of *C. sativus* were ground and homogenized in buffer (50 mM Tris-HCl pH7.5, 150 mM NaCl, 10 % glycerol, 2 mM EDTA, 5 mM DTT,

1 % IGEPAL CA630, 0.5 mM PMSF, 1 mM Na2MoO4, 1 mM NaF) with Complete protease inhibitor cocktail (Roche) and Halt Phosphatase Inhibitor Cocktail (Thermo Fisher Scientific) at 100 times dilution. To remove any cell debris, the supernatant was collected after 15,000 ×g centrifugation for 5 min at 4 ˚C. Then samples were mixed with SDS-Laemmli buffer and boiled for 3 min at 95 ˚C in a heat block. Each sample was separated by 8-16 % gradient or 12 % SDS-PAGE. After gel electrophoresis, samples were transferred to a PVDF membrane using a semi-dry blotting system (Bio-Rad). Membranes were then blocked in TBS containing 0.1 % polyoxyethylene sorbitan monolaurate (nacalai tesque) and 5 % skim milk (FUJIFILM Wako Pure Chemical Corp.). Anti-HA-Peroxidase, High Affinity (12013819001, Roche) (1:2,000) was used to detect HA-tagged proteins. Anti phospho-p44/42 MAPK (T202/Y204) (4370L, Cell Signaling) (1:2,000) and anti-mouse IgG, HRP-Linked Whole Ab Sheep (NA931, GE Healthcare) (1:10,000) were used to detect phosphorylated mitogen-activated protein kinases (MPKs) from *C. sativus* as a primary antibody and a secondary antibody, respectively. Anti-GFP from mouse $IgG_{1}\kappa$ (clones 7.1 and 13.1) (11814460001, Roche) (1:2,000) and anti-mouse IgG HRP-Linked Whole Ab Sheep were used to detect GFP-fused proteins as a primary antibody and a secondary antibody, respectively. For phosphorylated MPKs detection, PVDF Blocking Reagent for Can Get Signal (TOYOBO) and Can Get Signal Immunoreaction Enhancer Solution (TOYOBO) were used according to the manufacturer's protocol. Chemiluminescence was induced by SuperSignal West Femto Maximum Sensitivity Substrate (Thermo Fisher Scientific) and images were acquired by an ImageQuant LAS 4000 system (GE Healthcare).

**Method S5 Recombinant protein expression and purification**

Plasmids encoding SRN1 $^{H63A/\Delta SP}$-myc-6×His (pNK144) or SRN1 $^{H63A/H114A/\Delta SP}$-myc-6×His (pNK146) fused with α-factor secretion signal from *Saccharomyces cerevisiae* at N-termini under the control of *AOX1* promoter, which can be activated by methanol, were linearized by PmeI restriction enzyme and transformed into the *Pichia pastoris* SMD1163 strain according to manufacturer's protocol (Thermo Fischer Scientific). *P. pastoris* transformants were selected on YPD agar media with Zeocin at 100 μg $ml^{-1}$. To identify colonies highly expressing recombinant proteins, 8 transformants of each plasmid were grown in 5 ml buffered glycerol-complex medium (BMGY; 1% yeast extract, 2% peptone, 100 mM potassium phosphate pH6.0, 1.34% yeast nitrogen base with ammonium sulfate without amino acids, $4 \times 10^{-5}$% biotin, 1% glycerol) at 30 ˚C in a shaking incubator (300 rpm) for 16-24 hours. Cultured cells were harvested by centrifuging at 3000×g for 5 min at room temperature. The supernatants were discarded and cell pellets were resuspended in 5 ml buffered methanol-complex medium (BMMY; 1% yeast extract, 2% peptone, 100 mM potassium phosphate pH6.0, 1.34% yeast nitrogen base with ammonium sulfate without amino acids, $4 \times 10^{-5}$% biotin, 0.5% methanol) to an $OD_{600}$ of 1. To induce expression of recombinant proteins, resuspended cells were cultured in a shaking incubator (300 rpm) at 30 ˚C for 16 hours. Then cultures were centrifuged at 3,000×g for 5 min at 4 ˚C and supernatants were collected. To confirm expression of recombinant proteins, supernatants were separated by 4-15% poly-acrylamide gel electrophoresis using Criterion Tris-HCl precast gels (Bio-Rad). Gels were stained with Instant*Blue* (Sigma-Aldrich) following the manufacturer's protocol. For protein purification, *P. pastoris* transformants were cultured in 100 ml BMGY in 500 ml flasks. Cultured cells were harvested by centrifuging at 3000×g for 5 minutes at room temperature. Supernatants were discarded and cell pellets were resuspended in 200 ml BMMY to an $OD_{600}$ of 1.0 in 2 l baffled flasks and cultured in a shaking incubator (300 rpm) at 30 ˚C for 12-20 hours. A 50-ml aliquot of cultures were centrifuged at 5,000×g for 5 min to remove cells, then the supernatants were filtered through a syringe filter (GD/X 25 mm Syringe Filter, 0.45 µm; GE healthcare). These clarified supernatants were mixed with equal volume of 100 mM Tris-HCl, pH7.5 and 1 M NaCl and incubated with Ni sepharose excel (GE healthcare) for 1 hour. After that, the resin was washed with 20 mM Tris-HCl pH 7.5, 500 mM NaCl, and 30 mM imidazole. The bound protein was eluted with 20 mM Tris-HCl, pH7.5, 500 mM NaCl and 500 mM imidazole, and then concentrated and buffer exchanged to 20 mM Tris-HCl pH 7.5, 150 mM NaCl and 1 mM DTT using an ultrafiltration unit (Vivaspin turbo 15, 10 MWCO; Sartorius). Concentration of recombinant proteins was calculated using absorbance at 280 nm measured by a microvolume spectrophotometer, NanoDrop One (Thermo Fisher Scientific). Molar concentration of each recombinant proteins was calculated using molecular weight ($SRN1^{H63A/\Delta SP}$-myc-6×His; 14,396.99 Da, $SRN1^{H63A/H114A/\Delta SP}$-myc-6×His; 14,770.93 Da).

**Method S6** Linker ligation of RNAs

Linker DNA oligonucleotides (5'-ATCGTATCGTAGATCGGAAGAGCACACGTCTGAA-3') were chemically synthesized with 5' phosphorylation and addition of 2′,3′-dideoxycytidine at their 3′ end (Table S4) (Mito *et al.*, 2020). The 5'-end of linker DNA oligonucleotides was prepared as previously described (Mito *et al.*, 2020). Cleaved fragments of ssRNA2 on urea gel were excised and extracted using Oligo Clean & Concentrator kit (Zymo Research) according to manufacturer's protocol. ssRNA2 and the cleaved fragments were resuspended in 10 mM Tris-HCl pH7.0 and denatured at 95 °C for 2 min and then placed on ice for 3 min. ssRNAs were dephosphorylated using T4 polynucleotide kinase (New England Biolabs) as previously described (Mito *et al.*, 2020). Then, pre-adenylated linkers were added and treated with T4 RNA ligase 2 truncated K227Q (New England Biolabs) as previously described (Mito *et al.*, 2020). The reaction was separated using 15% urea gel in the same method as described in in vitro RNase assay section.
